# Sex-biased gene expression and gene-regulatory networks of sex-biased adverse event drug targets and drug metabolism genes

**DOI:** 10.1101/2023.05.23.541950

**Authors:** Jennifer L. Fisher, Amanda D. Clark, Emma F. Jones, Brittany N. Lasseigne

## Abstract

**Background:** Previous pharmacovigilance studies and a retroactive review of cancer clinical trial studies identified that women were more likely to experience drug adverse events (i.e., any unintended effects of medication), and men were more likely to experience adverse events that resulted in hospitalization or death. These sex-biased adverse events (SBAEs) are due to many factors not entirely understood, including differences in body mass, hormones, pharmacokinetics, and liver drug metabolism enzymes and transporters.

**Methods:** We first identified drugs associated with SBAEs from the FDA Adverse Event Reporting System (FAERS) database. Next, we evaluated sex-specific gene expression of the known drug targets and metabolism enzymes for those SBAE-associated drugs. We also constructed sex-specific tissue gene-regulatory networks to determine if these known drug targets and metabolism enzymes from the SBAE-associated drugs had sex-specific gene-regulatory network properties and predicted regulatory relationships.

**Results:** We identified liver-specific gene-regulatory differences for drug metabolism genes between males and females, which could explain observed sex differences in pharmacokinetics and pharmacodynamics. In addition, we found that ∼85% of SBAE-associated drug targets had sex-biased gene expression or were core genes of sex- and tissue-specific network communities, significantly higher than randomly selected drug targets. Lastly, we provide the sex-biased drug-adverse event pairs, drug targets, and drug metabolism enzymes as a resource for the research community.

**Conclusions:** Overall, we provide evidence that many SBAEs are associated with drug targets and drug metabolism genes that are differentially expressed and regulated between males and females. These SBAE-associated drug metabolism enzymes and drug targets may be useful for future studies seeking to explain or predict SBAEs.

## Background

In the U.S., adverse events (defined by the U.S. Food and Drug Administration [FDA] as any undesirable experience associated with using a medical product) resulted in an annually estimated 1.3 billion emergency room visits and an approximate 3.5 billion dollar economic impact [1–3]. Adverse events that are more likely to occur in one sex are called sex-biased adverse events (SBAEs) [4]. In 2001, the FDA removed ten drugs from the market; eight had female-biased adverse events [5]. Since then, several studies have found that women are twice as likely to experience an adverse event than men, based on adverse event case reports from the FDA Adverse Event Reporting System (FAERS) and World Health Organization (WHO) VigiBase database [6–10]. A retroactive review of cancer clinical trial studies found that women were more likely to experience adverse events from chemotherapy and immunotherapies, indicating that adverse event reporting bias is not necessarily the sole cause of SBAEs [9]. On the other hand, a VigiBase study found that men were more likely to experience adverse events that resulted in hospitalization or death [7]. Furthermore, the gender gap between men and women in the number of female-biased SBAEs increased during the coronavirus disease 2019 (COVID-19) pandemic [8].

While there are multiple sources of evidence for SBAEs, the biological differences that might result in SBAEs are still being investigated. Proposed causes of SBAEs include, but are not limited to, sex differences in body mass, hormones, pharmacokinetics, and liver drug metabolism enzymes and transporters [4]. One early hypothesis was that body mass differences between males and females resulted in SBAEs. While body mass differences are a known factor for drug response outcomes, many SBAEs are not explained by body mass differences [11,12]. Sex hormone differences have also been hypothesized to cause SBAEs and found to compete for drug transporters, compete with and inhibit enzymes, alter transcription, and interact with receptors on target cells [13]. Another previously investigated drug response factor is pharmacodynamics and pharmacokinetics [11]. Multiple studies have found sex differences in several pharmacokinetic metrics, such as the area under the curve of the plasma concentration of a drug versus time after dose or peak/maximum concentration [11]. Other studies have shown that many drug metabolism enzymes have differential gene expression and protein activity in male and female liver tissue [14–16]. Additionally, Oliva, et al. showed that in the Genotype-Tissue Expression (GTEx) project (n = 16,245 RNA-sequencing samples across 44 human tissues), 37% of genes had sex-biased tissue-specific expression and that these genes were enriched in drug metabolism gene sets (i.e., Gene Ontology [GO] and Kyoto Encyclopedia of Genes and Genomes [KEGG] terms: “Xenobiotic Metabolism, Mitochondrial Genes and Fatty Acid Oxidation,” “ Cellular Response to Hormones and Drugs,” “Drug Interaction and Response,” and “Drug Interaction”) [17].

Other studies have shown that regulatory relationships between transcription factors and genes differ between males and females in a tissue-specific manner that gene expression profiles alone can not identify [18,19]. Additionally, other studies have applied gene-regulatory networks to investigate drug mechanisms and predict drug sensitivity, repurposing candidates, and precision medicine [20–22]. In regards to sex differences in gene-regulatory networks, a previous study in colon cancer patients applied gene-regulatory network construction to colon cancer gene expression profiles from The Cancer Genome Atlas (TCGA) project and found that drug metabolism genes were more targeted in the female network than the male network in tumor tissue [23]. Sex-specific gene-regulatory networks of breast tissue have also shown sex differences in network communities (i.e., groups of transcription factors and genes that are more interconnected than other genes in the network) involved in different pathways, such as developmental and signaling pathways [18]. These sex differences in gene-regulatory networks across tissues could be potential factors in SBAEs.

We hypothesized that drug metabolism enzymes and drug targets of drugs associated with SBAEs had sex-biased gene-regulatory network properties compared to other genes and drug targets (**Figure 1**). First, we identified 416 drugs associated with SBAEs (i.e., SBAE-associated drugs) from cases reported in the FAERS database. We found that 32 drug metabolism enzymes and 84 drug targets were more likely to be targets of these SBAE-associated drugs than non-SBAE-associated drugs. KEGG-annotated drug metabolism genes also had sex differences in their liver gene-regulatory network neighborhoods and individual transcription factor gene relationships. For the 84 SBAE-associated drug targets, we found that their gene expression was more likely to be sex-biased and for those genes to be core genes (i.e., essential genes for gene-regulatory network community identification). These results support our hypothesis that the known drug metabolism enzymes and drug targets of drugs associated with SBAE have sex differences in gene expression and gene-regulatory networks.

**Figure 1:**
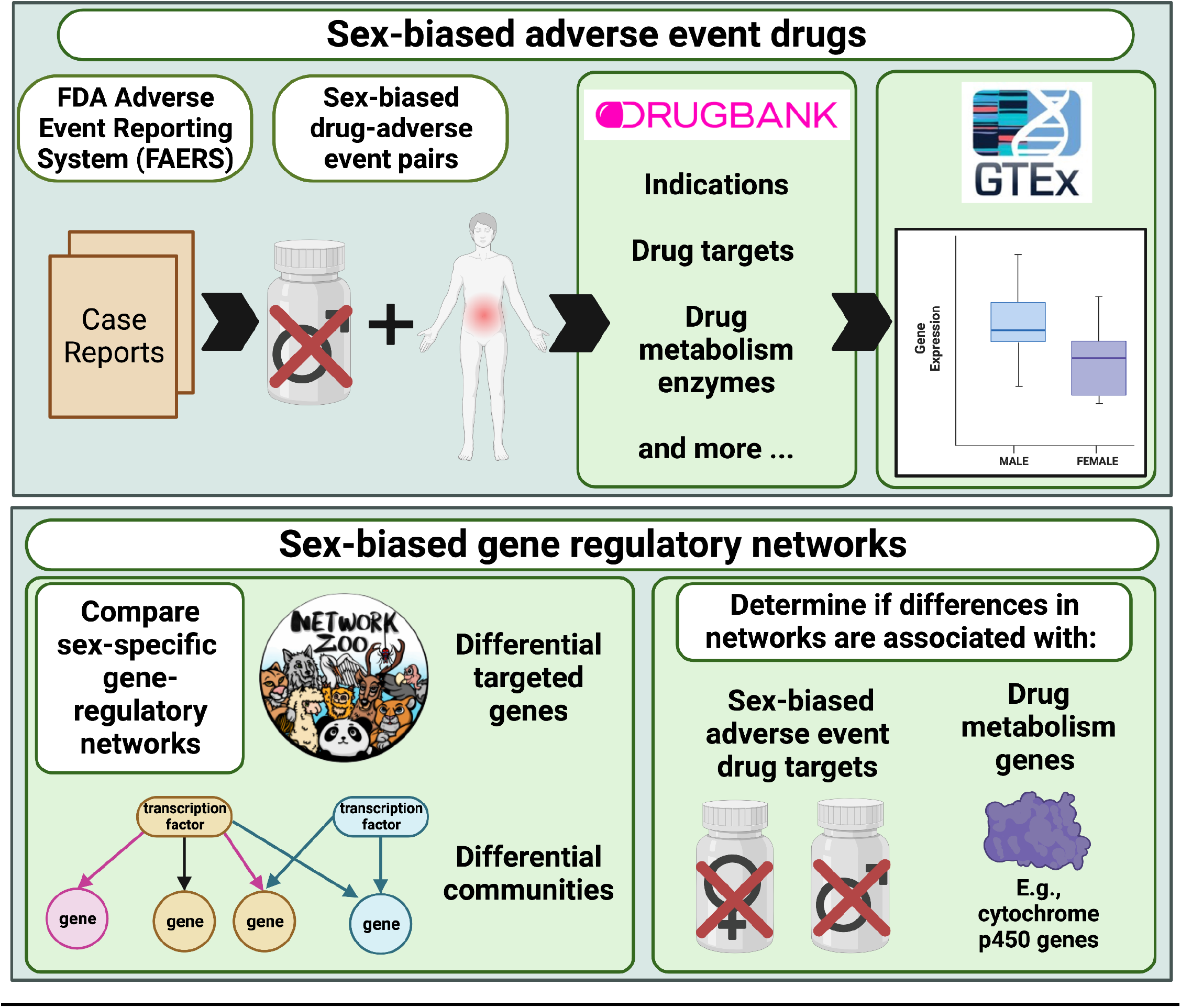
Graphical abstract of the study.

## Methods

### Scripts, dockers, and conda environment

The scripts for this project are available on Zenodo at https://zenodo.org/record/7938613#.ZGKFpdbMIaQ. In addition to the scripts there, the Docker images used for this analysis are publicly available on Docker Hub (jenfisher7/rstudio_sex_bias_drugs) and Zenodo (https://zenodo.org/record/7941598#.ZGOw5tbMKJE) (R version 4.2.2). For Fisher’s exact test calculations, we used a conda environment on the University of Alabama’s high-performance system, Cheaha, in an array format (SR_TAU_CELL_environment.yml) (R version 4.0.5).

Comprehensive computer environment and package version information is included in **Supplemental File 1**.

### Data download and exploration

#### Medical Dictionary for Regulatory Activities (MedDRA)

We downloaded the MedDRA database in July 2022 (Version 25.0) [24]. This database contains a 5-level hierarchy of medical terminology from lowest-level terms (e.g., “abnormal EEG”) to preferred terms (e.g., “electroencephalogram abnormal”), and finally to their highest-level terms, the system organ class terms (e.g., “investigations”). FAERS case reports contain MedDRA lower-level and preferred terms to annotate adverse events. We mapped all the lower-level terms to the system organ class terms to investigate groups of preferred terms and system organ class terms in the context of SBAEs.

The MedDRA database is a subscription database. We provide the workflow we used to format the downloaded database to the mappings used for the rest of this study in the provided scripts above.

#### FAERS

We downloaded curated FDA Adverse Event Reporting System (FAERS) data from the Zhang et al. study [8]. The FAERS database contains case reports of reported adverse events. We filtered the curated cases by country (i.e., “US”), qualification of the reporting party (i.e., “1” = physician, “2” = pharmacist, “3” = other health professionals), and cases that contained information for both sex (i.e., “gender”) and drugs. We also converted any lower-level MedDRA adverse event terms to their preferred terms via the MedDRA mappings described above. We only retained the newest case report for the three duplicated cases we identified.

#### GTEx

We downloaded preprocessed RNA-Seq count tables and metadata for the GTEx project with the Recount3 R package (version 1.8.0; accessed March 2022). We removed the sample GTEX-11 ILO, identified in previous literature as an individual who completed a gender-affirming surgery [25]. We also filtered samples to only those sequenced with the TRUSeq.v1 chemistry. We removed the following tissues due to being a sex-specific tissue or there being less than five samples in one sex: cervix-ectocervix, cervix-endocervix, fallopian tube, kidney-medulla, ovary, uterus, vagina, prostate, and testis. We also removed samples with a RIN score less than or equal to five. We recorded the final number of samples by sex for each tissue in **Supplemental File 2**.

In addition, we used the base R prcomp function to perform a principal component analysis (PCA) of the GTEx liver tissue samples to determine potentially confounding variables (i.e., age, RIN score, and ischemic time) that may affect gene expression and downstream results. We transformed the gene counts via DESeq2 (version 1.38.2) variance stabilizing transformation (vst). Then, we visually inspected PCA one through five, which had a cumulative percent variation of 41.4%, for clustering of samples based on age, RIN score, and ischemic time. We found samples clustered by ischemic time and RIN score with PC 1 (19.09% of variance explained) and PC 2 (8.916% of variance explained). These had a negative Spearman correlation with one another (rho= −0.5652864).

The methodology used to construct gene-regulatory networks, Passing Attributes between Networks for Data Assimilation (PANDA), required gene expression profiles. Before normalizing the GTEx data above, we filtered out genes with less than one count per million across all samples. We normalized the gene expression counts with a quantile shrinkage normalization via the R package YARN qsmooth function (version 1.24.0) [25] to remove variation due to technical variables in an unsupervised manner but with the “group” parameter to normalize the filtered counts data in a sex-aware manner to maintain the biological signal related to sample sex [25].

#### Protein-protein interaction and transcription factor motif information for gene-regulatory networks

The gene-regulatory network analysis required protein-protein interaction, transcription motif information, and RNA-Seq profiles. We downloaded the human protein-protein interaction network from the STRING database (accessed date: Jan. 2023; version 11.5). In prior studies, researchers applied protein-protein interaction networks to investigate the downstream effects of drugs and predict drugs for diseases as well as adverse events [26–28]. We adjusted the STRING network of known and predicted protein-protein interactions, including direct and indirect associations, for downstream gene-regulatory network construction by transforming edge weights to be between zero and one and focusing on highly confident interactions. To achieve this data transformation, we divided the raw interaction scores by 1000 and filtered for interactions greater than 0.7. We converted protein Ensembl IDs to HGNC gene symbols with the protein metadata in STRING. Additionally, we downloaded preprocessed transcription factor motif mapping (accessed date: Jan. 2023) [29]. We adjusted the TF-to-motif mappings to TF-to-gene mappings by mapping motifs to target genes to generate TF-gene-regulatory networks.

#### DrugBank

We downloaded DrugBank’s complete database (Version 1.5.8) in Jan. 2021. This database contains information about drugs, including drug approval, drug targets, drug metabolism genes, and indications. We also used and modified functions to access drug metabolism enzyme information from the drugbankR R package (version 1.5) to identify drug targets and metabolism enzymes of all drugs in the FAERS dataset.

### Fisher’s exact test and reporting odds ratio of FAERS database

To determine if drug and adverse event pairs were more likely to occur in one sex, we conducted a Fisher’s exact test and calculated the reporting odds ratio (ROR). We constructed contingency tables for each drug-adverse event combination for females compared to males. These tables included the following groups: A = the number of female patients with target drug-adverse event pairs, B = the number of female patients with the drug but not the same adverse event, C = the number of male patients with target drug-adverse event pairs, and D = the number of male patients with the drug but not the same adverse event. In addition, based on these contingency tables, we filtered to focus on more commonly used drugs-adverse event pairs in both sexes via the following thresholds:

- 30 cases of the drug-adverse event pair in at least one sex (A >= 30 or C >= 30)
- 50 cases of drug-adverse event pair across both sexes (A+C >= 50)
- 1000 cases for a drug across all four groups (A+B+C+D >= 1000)
- More than five adverse events in both sexes (A > 5 and C > 5).

Our approach was similar to the previous Yu et al. 2016 study [6]. We used the Beniamini-Hochberg (BH) procedure to adjust for multiple hypothesis testing. The calculated ROR from the Fisher’s exact test described if females (i.e., positive ROR) or males (i.e., negative ROR) were more likely to report the drug-adverse event combination. These ROR values were log-base2 transformed (i.e., logROR). A threshold of absolute logROR greater than one and the BH-adjusted p-value less than 0.05 identified sex-biased drug-adverse event pairs in the FAERS database. We determined if there were shared or different drugs and adverse events between the male- and female-biased drug-adverse event pairs via hypergeometric test for both drugs and adverse events.

### Identifying SBAE-associated drug target and drug metabolism genes

We annotated the 416 drugs identified as associated with an SBAE (i.e., SBAE-associated drugs) with their drug targets and drug metabolism enzyme genes from DrugBank [30]. To evaluate if drug targets were enriched in the drug target list of the 416 SBAE-associated drugs compared to randomly selected drugs, we performed permutation testing for all the drug targets of drugs in the FAERS database. We randomly selected the same number of drugs (i.e., 416) from FAERS and identified the number of drugs with that target, repeating the process 1,000 times. We performed a one-sample Wilcoxon signed rank test to determine if the number of SBAE-associated drugs with that drug target was higher than those for the randomly selected drugs. We applied a BH p-value adjustment. We filtered our top SBAE-associated drug targets by BH-p-value < 0.001. To focus our downstream analysis on SBAE-associated drug targets across SBAE-associated drugs, five SBAE-associated drugs had to have that known drug target. We also applied these same processing and filtering methods to drug metabolism genes.

### Sex-specific tissue gene-regulatory network construction and sex-specific community analysis

We applied the PANDA methodology to construct sex-specific gene-regulatory networks by tissue. PANDA integrates regulatory (TF Motif mappings from DNA-motifs curated by the Glass Lab [29]), protein-protein interaction (STRING Database [31]), and qsmooth normalized gene expression profiles from one sex and tissue via the pandaR package (version 1.30.0) [25]. We compared each tissue’s sex-specific network community structures via ALtered Partitions Across Community Architectures (ALPACA)’s differential modularity functionality (netZooR version 1.2.1). This algorithm determines if genes and transcription factors are differentially connected between networks, in this case, between the female and male networks. To determine the sex-specific communities, we used this approach to calculate the differential modularity score for each node (i.e., genes and transcription factors) in the sex-specific network, which was then used by the Louvian algorithm to assign a community for each node. This differential modularity score describes how important a node was to the resulting community assignment. We made the comparison twice: once to determine female-specific communities and once to determine male-specific communities. To determine female-specific communities, we assigned the female network as the perturbed network and the male as the baseline network. Then, to identify male-specific communities, we assigned the male network as the perturbed network and the female network as the baseline network.

### Sex-specific community core genes

Based on the differential modularity scores from our ALPACA sex-specific community results, we identified the core genes for each sex-specific community for each GTEx tissue included in our study. For each sex-specific community in a tissue, we identified the 100 genes with the highest differential modularity scores as core genes for that tissue sex-specific community based on [18]. After identifying these core genes, we conducted functional enrichment analysis via gprofiler2 (version 0.2.1) to determine enriched pathways for each sex-specific community across the GTEx tissues [32]. We used gprofiler2 to conduct cumulative hypergeometric probability tests to determine if there is an enrichment of the core genes in gene sets from Gene Ontology (GO) [33], KEGG [34], Reactome [35], WikiPathways [36], TRANSFAC [37], miRTarBase [38], Human Protein Atlas [39], and The Comprehensive Resource of Mammalian Protein Complexes (CORUM) [40].

### Drug metabolism genes’ sex-specific gene-regulatory relationships

To determine the network property differences between male and female gene-regulatory networks around drug metabolism genes, we downloaded the Kyoto Encyclopedia of Genes and Genomes (KEGG)’s drug metabolism gene list (KEGG_DRUG_METABOLISM_CYTOCHROME_P450.v2022.1.Hs.gmt) from MSigDB via Gene Set Enrichment Analysis (GSEA)’s website in Jan 2023 (https://www.gsea-msigdb.org/gsea/msigdb/human/collections.jsp) [34,41,42].

#### Weighted in-degree difference

We used the default parameters for the calcDegree function from the pandaR package to calculate the weighted in-degree for each drug metabolism gene for each sex-specific tissue network. By using the default parameters, we did not transform or filter the Z-Scores from the PANDA networks, as previously reported [21,23]. Then we compared weighted in-degree between the sexes by conducting a Wilcoxon rank sum test with continuity correction applied across each tissue and then made a Bonferroni-corrected p-value adjustment [43]. We also calculated the median of the weighted in-degree difference between the male and female drug metabolism genes for each tissue.

#### Sex-biased edges and targeting

We examined if genes had different proportions of sex-biased edges (i.e., predicted transcription factor and gene regulatory relationship z-score > 2 and only present in one sex’s network) between the male and female liver gene-regulatory networks. We defined each targeted gene as sex-divergent (i.e., the proportion of sex-biased edges in the male- and female-biased directions is between 0.4 and 0.6), female-biased (i.e., the proportion of sex-biased edges in the female direction is greater than 0.6), or male-biased (the proportion of sex-biased edges in the male direction is greater than 0.6) as previously reported by Lopes-Ramos et al. [19].

#### Specific edges activator and repressor relationships

For the liver sex-specific gene-regulatory networks, we determined the predicted regulation of the transcription factor and targeted drug metabolism genes by finding the Pearson correlation between a given transcription factor gene’s qsmooth normalized expression and the targeted drug metabolism gene’s qsmooth normalized expression. The Pearson correlation has been applied to PANDA network regulatory relationships in Kuijer et al. [44]. If the Pearson correlation was significant after Benjamini-Hochberg (BH) p-value adjustment and positive, we defined these edges as activator edges. If the Pearson correlation was significant after BH p-value adjustment and negative, we described these edges as repressor edges. If the correlation was not significant, we defined these relationships as undefined. We applied this in a sex-specific manner where we only correlated the transcription factor and target gene’s expression by sex and in a non-sex-specific manner.

Once we identified these predicted regulatory relationships, we determined which relationships (i.e., edges) differed or were the same between the male and female liver gene-regulatory networks for drug metabolism genes. Then, we used functional enrichment analysis via gprofiler2 to determine enriched pathways and annotated gene sets for sex-specific relationships (i.e., male activators, male repressors, female activators, male repressors) [32]. We compared the enriched Biological Process GO terms across these sex-specific relationships by semantic similarity based on the method proposed by Wang 2007 [45], which considers the relationship of GO terms in the GO hierarchy via the GOSemSim R package (version 2.24.0) [45,46]. We used functions from rrvgo (version 1.10.0) (getGoTerm, loadOrgdb, getGoSize, reduceSimMatrix) to create GO term common parent terms from the enriched pathways based on the parent term in the GO hierarchy and the Wang semantic similarity [47]. We plotted the enriched pathway results in a heatmap and clustered them using ComplexHeatmap (version 2.14.0) [48].

To determine if there are more activator- or repressor-predicted gene-regulatory relationships impacting drug metabolism genes in sex-specific networks, we determined the difference in the sum of the activator edge weights and the sum of the repressor edge weights for each sex. If this value for a drug metabolism gene is positive, there are more activator edges than repressor edges. If this value for a drug metabolism gene is negative, then more predicted repressor relationships exist. We applied a Wilcoxon rank sum test with continuity correction to determine if there was a difference in the targeting relationships between male and female liver gene-regulatory networks of drug metabolism genes.

### Permutation testing of sex-biased gene expression and sex-biased core genes of SBAE-associated drug targets and core genes of sex-specific communities

In a previous study with the GTEx project, Oliva et al. identified sex-biased expressed genes for 44 different tissues [17]. We downloaded the sex-biased gene expression results in March 2023 from the GTEx portal (https://www.gtexportal.org/home/datasets). We assessed SBAE-associated drug targets and drug metabolism enzyme genes for sex-biased gene expression across each tissue (i.e., the GTEx sex-biased gene sets). We performed permutation testing by randomly selecting 84 drug targets (the same number of SBAE-associated drug targets we identified) 1,000 times with a one-sample Wilcoxon signed rank test and a BH-multiple hypothesis test correction. We applied the same approach for the drug metabolism genes but with 71 or 64 for drug metabolism genes expressed in the liver tissue or the liver gene-regulatory networks, respectively. We repeated this permutation approach with the core genes of the sex-specific gene-regulatory communities for both SBAE-associated drug targets and drug metabolism genes.

## Results

### SBAE-associated drugs are enriched for known drug metabolism genes and drug targets

We first used the FAERS database to identify drug-adverse event pairs more likely to occur in one sex by requiring that each drug-adverse event pair have at least five cases for both sexes to ensure selection for sex-biased and not just sex-specific adverse events (i.e., where the condition occurs in one sex such as prostate cancer). Similar to previous studies with the FAERS database [6,8,10], we identified more drug-adverse event pairs with a female bias than a male bias (2132, female; 748, male) (**Figure 2A and Supplemental File 3**). When we investigated the most common SBAEs based on the number of significant drug-adverse event pairs, malignant neoplasm progression, acute kidney injury, and death were the top three male-biased adverse events. The top female-biased adverse events were alopecia, urinary tract infection, and drug hypersensitivity (**Supplemental Figures 1 & 2**). Some of these adverse events are known to have higher incidents in one sex, for example, male-biased malignant neoplasm progression and female-biased alopecia and urinary tract infection [6,10]. In accordance with previous literature using FAERS and other independent databases and studies, we identified drugs for both male-biased and female-biased drug-adverse event pairs that were identified as sex-biased, such as warfarin [49], cholecalciferol [50], prednisone [51], methotrexate [52], and denosumab [53], which were in the top three male and female-biased adverse events (**Supplemental Figure 3 & 4**).

**Figure 2:**
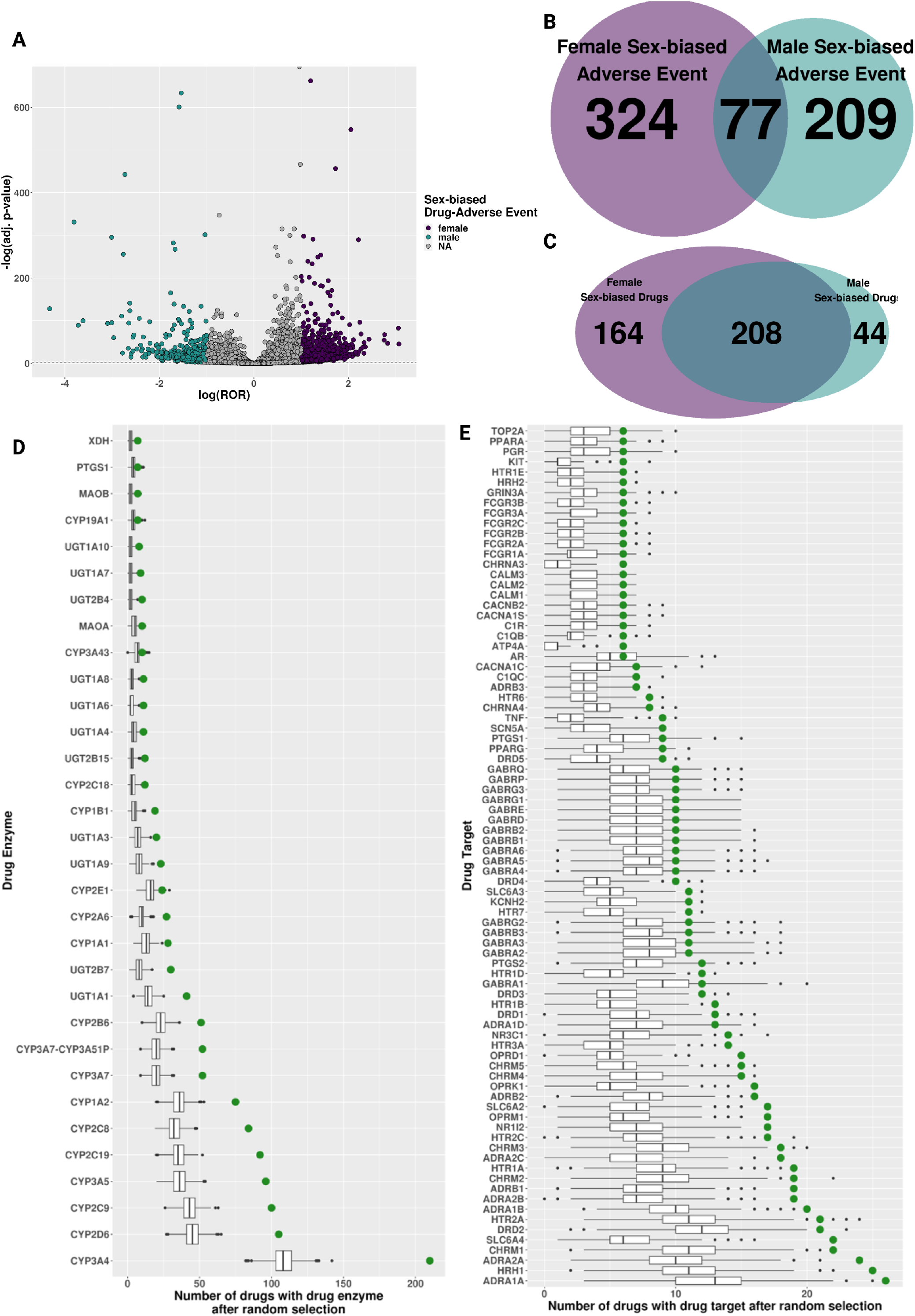
Sex-biased drug-adverse event pairs in FAERS. **A)** Volcano plot of the log-transformed reporting odds ratio (ROR) and the negative log-transformed Benjamini-Hochberg p-values from Fisher’s exact test of overlap of drug and SBAEs. **B)** Overlap of drugs from male- and female-biased drug-adverse event pairs. **C)** Overlap of adverse events from male- and female-biased drug-adverse event pairs. **D)** Boxplot of the number of randomly selected drugs with SBAE drug targets. **E)** Boxplot of the number of randomly selected drugs with SBAE drug enzyme. Green dot is the number of SBAE-associated drugs with drug target or drug enzyme.

Across the 2,880 significant sex-biased drug-adverse event pairs, we identified 610 male- or female-biased adverse events (i.e., the biological symptom). Of those, 77 adverse events were shared by male-biased and female-biased drug-adverse event pairs, which was not a significant overlap (p-value = 0.293085, hypergeometric test) (**Figure 2B**). However, there was a significant enrichment of shared SBAE-associated drugs between male-biased and female-biased drug-adverse event pairs. We found that 208 of the 416 unique SBAE-associated drugs from those drug-adverse event pairs were significantly associated with male and female drug-adverse event pairs (p-value= 2.524481e-11, hypergeometric test) (**Figure 2C)**. When we clustered the 50 most common drugs and the 50 most common adverse events from those 2,880 significant drug adverse-event pairs by their logROR, we found they clustered most strongly based on the SBAE (**Supplemental Figure 5**). This suggests that particular adverse events are more susceptible to SBAEs compared to particular drug mechanisms.

After identifying these sex-biased drug-adverse event pairs, we determined drug metabolism genes and targets enriched in the known drug metabolism enzymes and targets of SBAE-associated drugs via permutation testing of a matched number of random drugs. From this, we identified 32 known drug metabolism genes enriched in the known targets of SBAE-associated drugs (**Figure 2D)**. The most prevalent known drug metabolism enzyme across our identified SBAE-associated drugs, CYP3A4, is also the most prevalent drug metabolism enzyme for FDA-approved drugs overall, as an estimated 50% of FDA-approved drugs are metabolized by this enzyme [54]. For drug targets, we identified 84 drug targets enriched in the known drug targets of our identified SBAE-associated drugs compared to random drug selection (**Figure 2E)**. Two of the top three SBAE-associated drug targets were adrenergic receptors (i.e., *ADRA1A* & *ADRA2A*). These receptors are already known to have sex differences in locus coeruleus (LC)-norepinephrine (NE) arousal activity in different brain regions due to higher estradiol presence in females compared to males, which has been hypothesized to increase norepinephrine (NE) arousal in females [55].

### The gene-regulatory network neighborhoods around drug metabolism genes differ between males and females in the liver

Previous studies have identified that many drug metabolism genes have sex-biased gene expression in the liver [14–17], so we chose to investigate sex-specific liver gene-regulatory networks to determine if there are potential gene regulation differences of drug metabolism genes between males and females in the liver. Therefore, we constructed sex-specific liver gene-regulatory networks via the PANDA network methodology, representing the predicted regulatory relationship between transcription factors and target genes [56]. To investigate the immediate node neighbors (i.e., the network neighborhoods), we calculated the weighted in-degree (describes the magnitude of the predicted transcription factor regulation of a gene) of drug metabolism genes (as annotated by KEGG, n = 64) in the sex-specific liver networks [41]. We found that drug metabolism genes had a higher weighted in-degree (i.e., more targeted) in the female network compared to the male network (median of the degree difference of drug metabolism genes = 34.15683) (**Figure 3A**). To confirm if this relationship is specific to drug metabolism genes, we compared the degree difference of drug metabolism genes to other genes in these liver networks. We found a significant difference via a Wilcoxon rank sum test with continuity correction (W = 663550, p-value = 1.48 × 10^−8^) (**Figure 3A**). Additionally, we determined that this relationship is specific to only a few tissues by comparing male and female PANDA networks in 42 of the other GTEx tissues and applying the same Wilcoxon test followed by a Bonferroni correction (**Figure 3B**). Six other tissues besides the liver (“Brain Hypothalamus,” “Artery Coronary,” “Brain Anterior Cingulate Cortex [BA24],” “Heart Left Ventricle,” “Brain Nucleus Accumbens [basal ganglia],” and “Small Intestine-Terminal Ileum”) had more transcription factor targeting in female networks. In comparison, three other tissues (“Pancreas,” “Minor Salivary Gland,” and “Artery Aorta”) had more transcription factor targeting in male networks. However, the liver is the most significantly different tissue based on the Wilcoxon rank sum test with continuity correction with a Bonferroni correction (**Figure 3B**), and our results indicate that more transcription factors are predicted to regulate drug metabolism in the female liver gene-regulatory network compared to the male.

**Figure 3:**
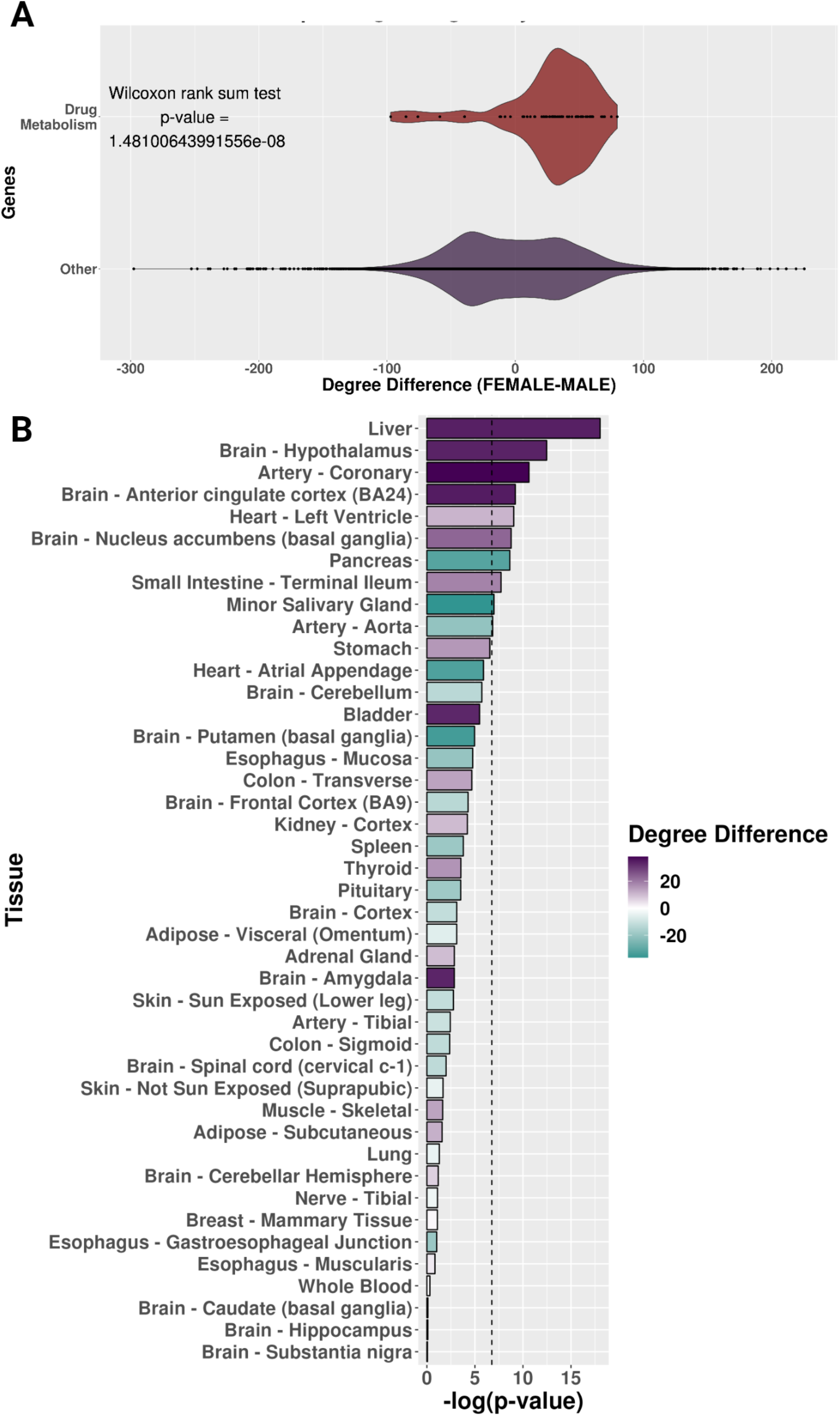
The neighborhood of sex-specific liver gene-regulatory networks around drug metabolism genes. **A)** Degree difference distributions of the sex-specific liver networks for drug metabolism genes compared to all other genes. **B)** Bar plot of the negative log-transformed p-value from the Wilcoxon test with colored bars showing the degree difference of drug metabolism genes between female and male sex-specific networks by tissue. For the bar plots, purple is a higher degree in female networks, and teal is a higher degree in male networks. The black dotted line is the p-value cutoff for the Bonferroni hypothesis test correction.

We were also interested in determining groups of transcription factors and genes that are more interconnected in their edge relationships in one sex-specific network than the other sex-specific network (i.e., sex-specific communities) in the liver. We identified eight female-specific communities and 11 male-specific communities. When we investigated which communities contained the most drug metabolism genes, we determined that female community number two included 41 of the 64 drug metabolism genes, and the other drug metabolism genes were contained in four other female-specific communities (**Supplemental Figure 6**). However, in the male network, we found that drug metabolism genes were divided between two male-specific communities (communities four and five), which contained 19 and 17 of the 64 drug metabolism genes, respectively (**Supplemental Figure 6**). The eight other male-specific communities contained the remaining 28 drug metabolism genes. We conducted functional enrichment analysis on the core genes (core genes have the highest differential modularity score, a score used to determine the sex-specific communities) of all the liver sex-specific communities. Several transcriptional pathways from KEGG and Reactome were similar between the female and male drug metabolism enzymes communities, such as the RUNX protein-associated pathways and nuclear receptor transcription pathway (i.e., liver female-specific community number two and male-specific communities numbers four and five). However, male-specific community number four had several pathways associated with FOXO transcription activities (i.e., “FOXO-mediated transcription of cell death genes,” “FOXO-mediated transcription,” “Regulation of FOXO transcriptional activity by acetylation,” “Regulation of localization of FOXO transcription factors,” and “FOXO-mediated transcription of cell cycle genes”) that were not shared with female-specific community number two and male-specific community five (i.e., the sex-specific liver communities with the most drug metabolism genes). Interestingly, previous aging studies have found that sex-dependent single nucleotide polymorphisms (SNPs) in *FOXO1A* (a protein important for regulating energy metabolism in the liver) are associated with increased longevity in females [57,58] (**Supplemental File 4**).

For each community, we found differing transcription factor and gene differential modularity scores, which describe how important that transcription factor or gene is to community integrity (i.e., if the node was removed, would the community structure still be present). We found that four drug metabolism genes, *CYP2E1*, *CYP3A43*, *GSTM4*, and *UGT2B17,* were also core genes in the male-specific liver communities (i.e., these genes had the highest differential modularity score). There was no significant enrichment of core genes in the drug metabolism gene set compared to randomly selected genes for sex-biased, female-biased, and male-biased liver core gene lists **(Supplemental Figure 7)**.

### Liver sex-specific networks have different drug metabolism gene-regulatory relationships

We next investigated the network edges representing the predicted regulatory relationship between a given transcription factor-gene pair that involved a drug metabolism gene (**Figure 4A**). Across all the known drug metabolism enzyme genes, we identified three female-biased targeted genes, *GSTO2*, *CYP2D6*, and *ALDH3A1* (i.e., the proportion of sex-biased edges in the female direction was greater than 0.6), and four male-biased targeted genes, *MAOA*, *AOX1*, *MGST1,* and *ALDH3B1* (i.e., the proportion of sex-biased edges in the male direction was greater than 0.6). When we compared the sex-biased targeted genes and the SBAE-associated drug metabolism enzymes, both *CYP2D6* and *MAOA* were drug metabolism genes of SBAE-associated drugs.

**Figure 4:**
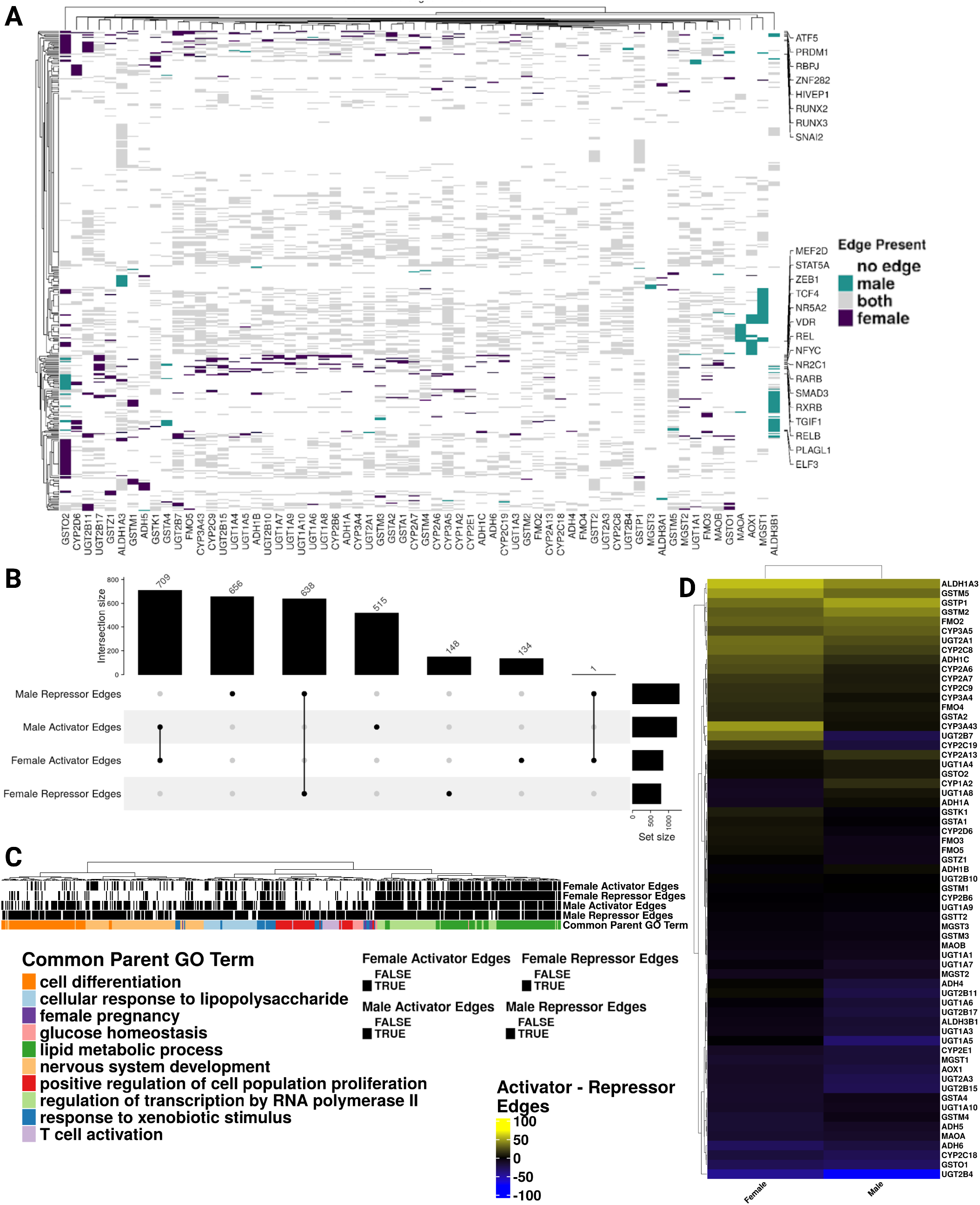
Predicted gene-regulatory relationships of drug metabolism genes in the sex-specific networks of the liver differ. **A)** Heatmap of the male and female associated liver gene-regulatory edges of drug metabolism genes (x-axis) and transcription factors (y-axis). The coloring on the heatmap indicates the network(s) where an edge was present. The y-axis is annotated with the transcription factors with the most male or female edges. **B)** Upset plot showing the intersection of activator and repressor transcription factor-gene edges in the female and male-specific liver networks. **C)** A dendrogram of the enriched GO Biological Process terms of unique female and male activator/repressor edges. The “Common Parent GO Term[s]” clustering and selection were based on Wang’s semantic similarity. **D)** A heatmap of the difference in activator and repressor edges of drug metabolism genes in the sex-specific liver networks (i.e., if the difference is positive, the drug metabolism gene is activator-targeted; if it is negative, it is repressor-targeted).

We calculated the Pearson correlation between transcription factors and drug metabolism genes and significance to identify types of potential regulatory relationships (i.e., BH p-value < 0.05). A significant positive correlation was an activator relationship, and a significant negative correlation was a repressor relationship. About 52% of these edge relationships were unique to one sex (**Figure 4B**). Interestingly, the only edge to have an opposite relationship between males and females was *ONECUT1*_*GSTM3*. While it was predicted to have an activator relationship in the female liver network (rho = 0.3511445), it was predicted to have a relatively weaker repressor relationship in the male liver network (rho = −0.1877166). Altogether, these results highlight that 1453 of the 2801 edges (∼51.87%) involving drug metabolism genes were unique to either sex. However, when the edges were present in both sexes, they had the same edge relationship (i.e., activator or repressor). Only 1 of the 1348 shared edges (∼0.07%) had an opposite relationship between males and females.

Given that about half of the drug metabolism gene edges were unique to one sex, we determined which GO Biological Processes were enriched between the distinctive male and female edges. We applied semantic similarity analysis to determine how similar the enriched GO terms were between the female and male activator and repressor edges and the common parent GO term of all the enriched GO terms (**Figure 4C** and **Supplemental File 5**). For all the unique sex-specific edges, “regulation of transcription by RNA polymerase II” (unsurprising given the abundance of transcription factors due to the examination of gene-regulatory relationships) and “lipid metabolic process” common parent GO terms were enriched. Male activator and repressor gene sets had more GO terms enriched for the following common parent GO terms than the female activator and repressor gene sets: “T-cell activation,” “nervous system development,” “glucose homeostasis,” “cellular response to lipopolysaccharide,” “cell differentiation,” “female pregnancy,” “positive regulation of cell population proliferation,” and “response to xenobiotic stimulus”. The parent term “female pregnancy,” which had more enriched GO terms with the male repressor and activator edges, included the following GO terms that relate to reproduction: “female courtship behavior,” “reproductive process,” “reproduction,” “developmental process involved in reproduction,” “multi-organism reproductive process,” and “female pregnancy” (**Supplemental File 5**). Overall, the number of male-biased common parent GO term groups (i.e., 8 of 10 common parent GO terms across all enriched pathways) indicates that liver male-specific edges of drug metabolism genes are potentially involved in more biological programs than female-specific edges.

For each drug metabolism enzyme gene, we determined if it had more activator or repressor targeting in the sex-specific networks (similar to the degree difference analysis in **Figure 3A**). In the female-specific liver network, we found 30 activator-targeted drug metabolism genes (i.e., a positive difference of activator and repressor edges of drug metabolism genes in the sex-specific liver networks) and 33 repressor-targeted drug metabolism genes (i.e., a negative difference of activator and repressor edges of drug metabolism genes in the sex-specific liver networks) (**Figure 4D**). However, in the male-specific liver network, we identified 24 activator-targeted and 39 repressor-targeted drug metabolism genes (**Figure 4D**). In addition, 16 drug metabolism genes had opposing targeting relationships between males and females. *GSTM1, UGT1A8, CYP1A2, ADH1B,* and *ADH1A* were more activator-targeted in the male liver network, while they were more repressor-targeted in the female liver network. However, *ADH4, CYP2C19, CYP2D6, FMO3, FMO5, GSTA1, GSTK1, GSTZ1, UGT2B10, UGT2B11,* and *UGT2B7* were more activator-targeted in the female liver network and more repressor-targeted in the male liver network. Five of the 11 differentially targeted drug metabolism genes were SBAE-associated drug metabolism enzyme genes, which might explain some SBAEs, including those associated with CYP2D6, which metabolizes 25% of drugs currently on the market (**Figure 2D**) [59]. However, we determined that the activator and repressor targeting of the drug metabolism genes in males and females was not significantly different (Wilcoxon rank sum test with continuity correction, W = 2347, p-value = 0.07735).

### SBAE-associated drug targets were more likely to have sex-biased gene expression and be core genes in sex-specific gene-regulatory networks than other drug targets

Building on previously identified sex-biased gene expression in GTEx, we determined if the SBAE-associated drug targets we identified above also had sex-biased gene expression (**Figure 5A**) [17]. We found that 52 of the 84 (∼ 62%) SBAE-associated drug targets had sex-biased gene expression in at least one tissue (29 had higher male gene expression in at least one tissue, and 35 had higher female gene expression in at least one tissue). We conducted permutation testing to determine if our identified SBAE-associated drug targets were more likely to have sex-biased gene expression than random sets of drug targets (**Supplemental Figure 8**). Across the GTEx tissues, we found that our identified SBAE-associated drug targets were significantly enriched (BH-adjusted p-value < 0.05) for sex-biased gene expression compared to randomly selected drug targets with respect to both female-biased (24 of 44 tissues) and male-biased (7 of 44 tissues) gene expression gene sets. The enrichment of SBAE-associated drug targets with sex-biased gene expression indicates that SBAEs could be due to drugs perturbing genes with sex-biased gene expression.

**Figure 5:**
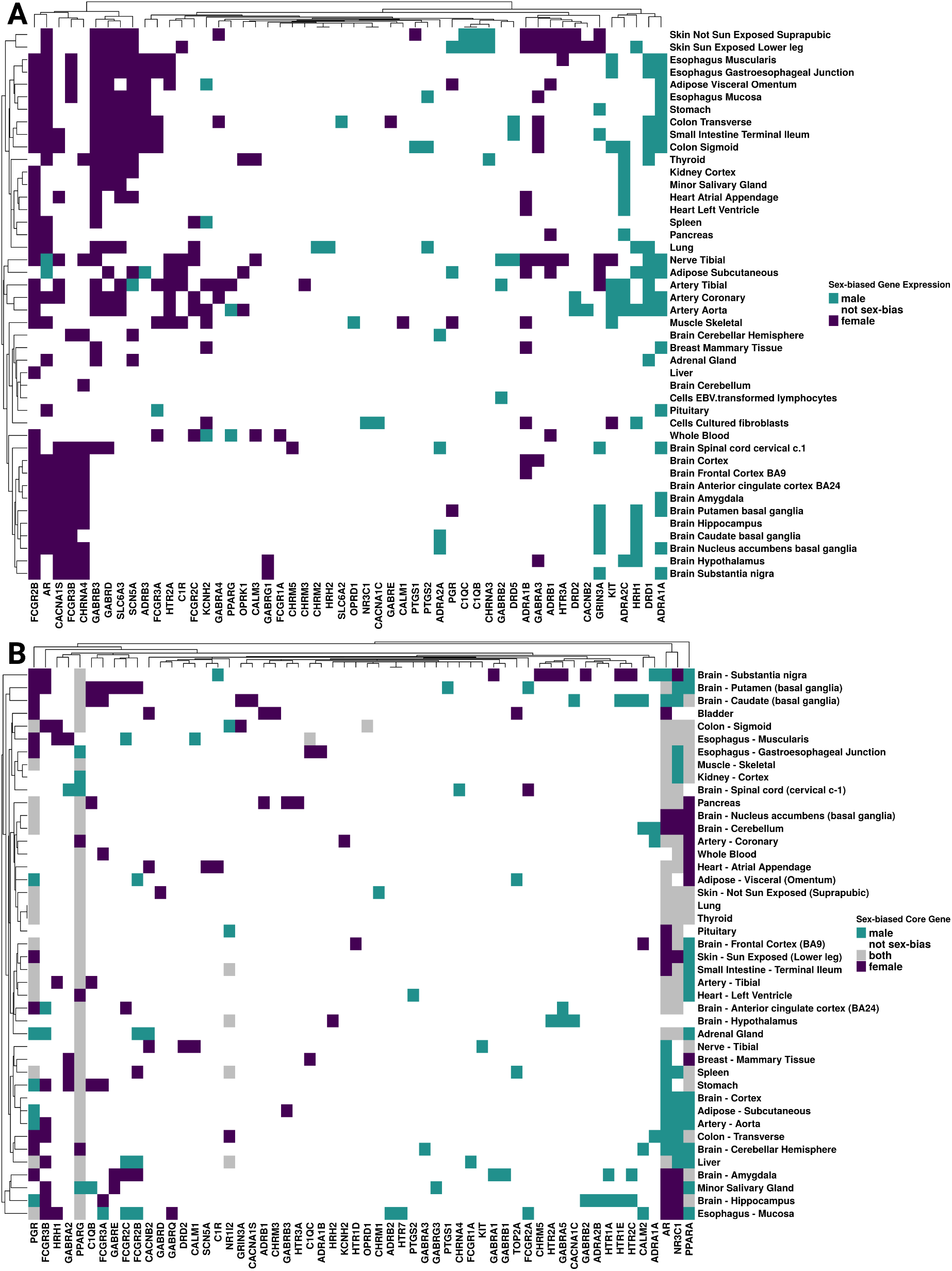
Most SBAE-associated drug targets have sex-biased gene expression or are sex-specific community core genes. **A)** Sex-biased gene expression of SBAE-associated drug targets across tissues. **B)** SBAE-associated drug targets were sex-specific gene-regulatory network core genes across tissues. Euclidean distance clustering with ComplexHeatmap’s complete algorithm was used for both heatmaps.

Furthermore, we explored *ADRA1A*, *DRD1*, and *ADRA2C,* the top three male-biased SBAE-associated drug targets based on the Euclidean distance clustering of sex-biased gene expression across tissues (**Figure 5A**) [48]. 26, 13, and 18 SBAE-associated drugs had *ADRA1A*, *DRD1*, and *ADRA2C* as a drug target, respectively. Both *ADRA1A* and *ADRA2C* are adrenergic receptors and are important for locus coeruleus (LC)-norepinephrine (NE) arousal activity [55]. While the adrenergic α2C-receptors (*ADRA2C*) have been shown to influence aggression behaviors in male mice, neither study investigated female mice in their study design [60,61]. Of the 52 male-biased drug-adverse event pairs with SBAE-associated drugs with *ADRA1A* as a drug target, 20 male-biased adverse events were related to psychiatric disorders (**Supplemental Figure 9A)**. In contrast, only one of the 126 female-biased *ADRA1A* adverse events was a psychiatric disorder (**Supplemental Figure 9B**). The highest male-biased adverse event preferred term was aggression which has the system organ class assignment of psychiatric disorder (**Supplemental Figure 9C**). *ADRA1A* had male-biased gene expression in five GTEx brain regions compared to females (“Brain Substantia Nigra,” “Brain Nucleus Accumbens Basal Ganglia,” “Brain Putamen Basal Ganglia,” “Brain Amygdala,” and “Brain Spinal Cord Cervical c.1”) (**Figure 5A**). Additionally, 18 SBAE-associated drugs had *ADRA2C* as a drug target. 17 of 31 male-biased drug-adverse event pairs of SBAE-associated drugs with *ADRA2C* as a drug target were male-biased for psychiatric disorder adverse events (**Supplemental Figure 10A)**. In contrast, only one of the 63 female-biased adverse events was a psychiatric disorder (**Supplemental Figure 10B**). Aggression was the highest preferred term for male-biased drug-adverse event pairs with *ADRA2C* as a drug target (**Supplemental Figure 10C**). This gene’s expression is only sex-biased in the hypothalamus (male-biased) (**Figure 5A**). Lastly, *DRD1*, a D1 dopamine receptor-coding gene, is the most abundant dopamine receptor subtype in the central nervous system [62]. 12 of the 21 male-biased drug-adverse event pairs of SBAE-associated drugs with *DRD1* as a drug target also had male-biased psychiatric disorder adverse events (**Supplemental Figure 11A)**. However, only one female-biased psychiatric disorder adverse event was associated with a drug that targets *DRD1* (**Supplemental Figure 11B**). However, *DRD1* does not have sex-biased gene expression in GTEx brain tissue RNA-Seq data (**Figure 5A**). Additionally, aggression had three male-biased adverse events, the highest along with hallucination, for SBAE-associated drugs with *DRD1* as a drug target (**Supplemental Figure 11C**). *DRD1* genetic variation and gene expression in different brain regions has been implicated with aggression, cognitive impulsivity, and psychosis [63–67]. It is currently unclear from the literature if there is documentation of sex-biased *DRD1* gene expression affecting aggressive phenotypes.

The top three female-biased drug targets with the highest number of tissues with female-biased gene expression were *FCGR2B*, *AR*, and *CACNA1S* [48]. 6, 28, and 6 SBAE-associated drugs had *FCGR2B*, *AR*, and *CACNA1S* as drug targets, respectively. 7 of the 45 female-biased drug-adverse event pairs with SBAE-associated drugs targeting *FCGR2B* had female-biased adverse events related to general disorders and administration site conditions, and six female-biased drug-adverse event pairs were infections and infestations (**Supplemental Figure 12**). *FCGR2B* is expressed by many immune cells, and previous literature identified that macrophages have higher *Fcgr2b* expression in female mice [68,69]. In addition, many SNPs of *FCGR2B* are associated with an increased risk of systemic lupus erythematosus, an autoimmune disease known to be more prevalent in females than males [69–71]. *AR* is the androgen receptor that interacts with androgens, including testosterone, and this receptor is important for sex differences in neural circuitry and metabolism [72]. Its expression was female-biased in 26 tissues, including 9 of 13 brain regions (**Figure 5A**). However, its expression was also male-biased in the tibial nerve and the subcutaneous adipose tissue (**Figure 5A**). While this gene’s expression was female-biased across 26 GTEx tissues, perturbing *AR* is associated with both male- and female-biased adverse events related to psychiatric disorders and nervous system adverse events, respectively (**Supplemental Figure 13A & B**). 11 of the 110 female drug-adverse event pairs we identified with *AR* as a drug target had female-biased nervous system adverse events, and 11 of the 60 male drug-adverse event pairs were related to male-biased psychiatric disorders (**Supplemental Figure 13A & B**). Lastly, *CACNA1S* has female-biased expression in 11 of the 13 GTEx brain regions (**Figure 5A**), and 9 of the 58 female-biased drug-adverse event pairs with *CACNA1S* as a drug target were related to the nervous system (the second highest number of female-biased drug-adverse events) (**Supplemental Figure 14**). Overall, these cases highlight a potential connection between a drug target’s sex-biased gene expression and their most common adverse events.

We also hypothesized that SBAEs might be due to drug targets that are core genes of sex-specific communities. We built sex-specific tissue networks for 43 different tissues based on the gene expression data from the GTEx project. We identified sex-specific communities within each tissue using the same process for the liver gene-regulatory network analysis above. We selected 100 core genes for each sex-specific community from the top 100 highest differential modularity score genes in each sex-specific community. In total, 58 of the 84 (∼ 69%) SBAE-associated drug targets were a core gene of a sex-specific gene-regulatory network in at least one tissue (**Figure 5B**). 41 of the 84 (∼ 49%) SBAE-associated drug targets were male core genes in at least one tissue, and 43 of the 84 (∼ 51%) drug targets were female core genes in at least one tissue. In total, SBAE-associated drug targets were enriched for female sex-specific community core genes in 34 tissues and for male sex-specific community core genes in 32 tissues (permutation testing with BH-p-value <0.05) (**Supplemental Figure 15**). We found that the nuclear receptors *PPARA*, *PGR*, *PPARG*, *AR*, and *NR3C1,* known to be regulated by hormone signaling (including sex hormones [73]), were drug targets and core genes to more than half of the tissues analyzed [73]. Additionally, *PPARA* is not differentially expressed between males and females in the GTEx dataset, while the other core genes had sex-biased gene expression in at least one tissue. We found one study where a PPARα agonist treatment before a stroke was neuroprotective in male mice but not females, suggesting that perturbation of PPARα might be sex-dependent [74]. Therefore, *PPARA* is an example of a gene that was a sex-biased gene-regulatory core gene without sex-biased gene expression, but where there was literature evidence of a sex-dependent PPARα agonist treatment drug response.

Overall, we found that 71 of the 84 (∼ 85%) drug targets had either sex-biased gene expression (∼ 62%) or high differential modularity in sex-specific communities (∼ 49%) (**Supplemental File 6**). 39 of 71 drug targets overlapped between the sex-biased gene expression list and the core gene list, and 13 and 19 drug targets were only identified by sex-biased gene expression or core genes, respectively. We found that 194 of the 389 (∼ 50%) SBAE-associated drugs we identified had at least one of the 71 drug targets we identified as having sex-biased gene expressions or core genes of sex-specific communities. These SBAE-associated drug target sex-biased differences in gene expression and gene-regulatory networks support our hypothesis that SBAE-associated drug targets are enriched for sex differences in gene expression.

## Discussion

Prior pharmacovigilance studies have identified SBAEs, and multiple studies have investigated sex differences in gene expression and gene-regulatory networks, implicating their role in drug metabolism and response [6–11,14,18,23]. In this study, we sought to identify if there are sex differences in gene expression and the gene-regulatory networks of the drug targets of SBAE-associated drugs. We first identified 32 drug metabolism enzymes and 84 drug targets enriched for SBAE-associated drugs. The liver gene-regulatory neighborhood and edges differed for drug metabolism enzymes between male and female gene-regulatory networks. Additionally, we found that SBAE-associated drug targets were more likely to be sex-biased expressed genes and the core genes of sex-specific gene-regulatory communities than randomly selected drug targets. Our findings support the hypothesis that some SBAEs may be due to drugs perturbing genes with sex-biased gene expression or gene-regulatory network properties.

Like previous pharmacovigilance studies with similar methods, we identified more than twice as many female SBAEs as male SBAEs [6,8,10]. Additionally, we identified 610 male- or female-biased adverse events, and the top three male- and female-biased drugs associated with SBAEs were also identified previously [49–53]. These findings are clinically relevant. For example, the second most common male-biased adverse event we identified was acute kidney injury (AKI). It is known that drug-induced AKI is a common cause of AKI, accounting for approximately 19% of AKI in hospital settings [75,76]. We also identified drug hypersensitivity as female-biased, supported by previous reviews of pharmacovigilance studies [77] and evidence suggesting multifactorial explanations for why females are more likely to have drug hypersensitivity, including potential genetic and epigenetic causes [78]. Our results highlight the need for more research investigating causal relationships between adverse events, drug mechanisms, and sex differences.

In this study, we also expanded upon previous studies examining sex differences in drug metabolism genes in the liver. CYP3A4 has higher gene expression, protein expression, and activity in females and was the most common drug metabolism enzyme across the SBAE-associated drugs we identified [79]. We also found that *CYP3A43* was a male core gene. In addition, two of the SBAE-associated drug targets, *CYP2D6* and *MAOA*, were female- and male-biased targeted in the sex-specific liver gene-regulatory networks. While *MAOA* was the 24th most common SBAE-associated drug metabolism gene, *CYP2D6* was the second most common SBAE-associated drug metabolism gene, and it was estimated 25% of drugs on the market use the CYP2D6 enzyme [59]. We identified two common drug metabolism enzymes enriched in SBAE-associated drugs with sex differences in liver gene-regulatory networks.

We further applied gene-regulatory network methodology to explore if there are predicted sex differences in gene regulation in the liver. We found that drug metabolism genes in the liver female gene-regulatory network are targeted more than in the male network via the weighted in-degree difference.In addition, the unique male liver network edges were involved in multiple biological processes, including “T-cell activation,” “nervous system development,” and “glucose homeostasis.” Our results support a potential female neighborhood structure where drug metabolism genes are more targeted than in the male liver network, and the transcription factors that regulate drug metabolism genes are involved in fewer biological programs than in the male network. This network structure further underscores that the influence of sex is complex for liver drug metabolism gene expression. For example, multiple biological processes utilize the same transcription factors as drug metabolism genes in the male-specific liver network. One common parent GO term from our GO semantic similarity analysis of the unique sex-specific edges of the drug metabolism genes was “glucose homeostasis.” Male repressor and activator edges were enriched for “glucose homeostasis,” “carbohydrate homeostasis,” “regulation of hormone levels,” “cellular glucose homeostasis,” “homeostasis of number of cells,” and “cellular response to glucose stimulus.” A prior study found that long-term exposure to abnormal glucose levels affected the activity of drug metabolism enzymes in primary hepatocytes, but unfortunately, it did not state the donor sex of the cells used in the experiment or discuss the impact of sex [80]. Future studies are needed to delineate the role of gene-regulatory sex differences in drug metabolism to determine the relative contribution of gene expression, protein expression, and protein activity to SBAE.

In this study, we sought to evaluate our hypothesis that drugs associated with SBAEs are perturbing sex-biased gene expression and gene-regulatory networks. The 84 enriched SBAE-associated drug target genes we identified were more likely to be expressed in a sex-biased manner and to be core genes than other drug targets across several tissues. In addition, we found that some of the SBAE-associated drug targets with sex-biased gene expression were associated with a common adverse event and associated sex-biased gene expression in the tissue manifesting that adverse event. For example, *ADRA1A* and *ADRA2C* were both associated with the male-biased adverse event of aggression. We found that these genes have higher expression in many brain regions in males than in females. However, we identified other SBAE-associated drug targets with more complicated expression and network patterns. For example, *AR,* which codes for the androgen receptor, is known to be influenced by the sex hormone testosterone, which is dynamically secreted over hours and decades [81]*. AR* was among the top three female-biased expressed genes and sex-biased core genes in all 43 tissues **(Figure 5**). We found that it had common female-biased adverse events of nervous system disorders and male-biased adverse events of psychiatric disorders, suggesting *AR* may contribute to different SBAEs for each sex.

There are some limitations to the current study. First, these results are associations and not causal relationships, so future studies are needed to investigate their causality. We have provided our computational workflows and complete results to facilitate their use as a resource for future studies. Also, we acknowledge that FAERS is based on case reports susceptible to reporting biases and missing data, and GTEx samples are skewed toward male subjects and older individuals, with ∼79% of the subjects at age 40 or above. Additionally, we found that RIN score and ischemic time are potentially confounding variables in the GTEx gene expression data. This technical variation could not be incorporated into the PANDA methodology. Still, the sex-biased gene sets from Oliva et al. included RIN score and ischemic time as covariates in their analyses [17]. As we previously reviewed [4], other pharmacokinetic, pharmacodynamic, biological, socioeconomic, and environmental factors can influence drug response in combination with sex, such as solubility of the drug, body fat percentage, diet, ancestry, and age. Additionally, with respect to age, it has been noted in the literature that males experience more adverse events than females before puberty, but this changes after puberty [4]. Our study used the GTEx dataset in which ∼79% of the samples are 40 and above; therefore, we were limited in investigating SBAEs that occur before puberty. This highlights another study limitation, since menopause occurs at an average age of around 50 and therefore, many of our female GTEx samples are likely post-menopausal (though this was not reported). There are many hormonal and gene expression changes that occur during and after menopause, which may be related to the occurrence of sex-biased adverse events and could be followed up in future studies [82]. Other potential complexities of SBAEs could be due to drug delivery and/or dosing. That information is limited in the FAERS dataset due to lack of reporting. Additionally, a drug could disrupt one tissue which then affects multiple other tissues or multiple tissues could be impacted by a drug directly. These potential complexities could be further explored in future studies.

Lastly, one critical limitation of this study is that we relied on bulk tissue RNA-Seq profiles from GTEx. With the invention of single-cell and spatial sequencing technology, the research community has determined gene expression differences between cell populations within a tissue sample [83–85]. For example, single-cell technology was recently used to detect sex-biased gene expression and gene-regulatory networks in mouse brain and heart tissues [86]. Concerning our study, for example, because *FCGR2B* is associated with the immune system, which was difficult to investigate with bulk gene expression profiles, other literature sources were added to help clarify our association of female-biased infections and infestations adverse events and FCGR2B female-bias expression in immune cells [68–71]. Another limitation of bulk tissue profiles was highlighted by tissues with multiple tissue regions sequenced by GTEx, like the brain. Because of this, we could associate more neurological and psychiatric disorder adverse events with drug targets than other tissues and adverse events. Therefore, future studies using a similar methodology applied to additional tissue subregions and single-cell profiles, particularly for organs involved in drug metabolism like the kidney and liver, are critical.

## Conclusions

Here, we used data mining and network approaches to investigate not only the gene expression of both drug metabolism genes and drug targets of drugs associated with SBAEs but also examined network properties of sex- and tissue-specific gene-regulatory networks. While previous studies have focused on drug metabolism enzymes, we also investigated the known drug targets of SBAE-associated drugs. Overall, we found supporting evidence that SBAEs could be caused in part by sex differences in drug metabolism enzyme and drug target gene expression and gene-regulatory network properties. These results are a valuable resource for future studies determining SBAE mechanisms to predict and prevent SBAEs and for sex-aware drug development and repurposing.

## Abbreviations

SBAEs: Sex-biased adverse events
FDA: U.S. Food and Drug Administration
FAERS: FDA’s Adverse Event Reporting System
WHO: World Health Organization
COVID-19: coronavirus disease of 2019
GTEx: Genotype-Tissue Expression project
TCGA: The Cancer Genome Atlas
GO: Gene Ontology
KEGG: Kyoto Encyclopedia of Genes and Genomes
MedDRA: Medical Dictionary for Regulatory Activities
PANDA: Passing Attributes between Networks for Data Assimilation
ALPACA: ALtered Partitions Across Community Architectures
PCA: principal component analysis
ROR: reporting odds ratio
BH: Benjamini-Hochberg
logROR: log reporting odds ratio
SNPs: Single nucleotide polymorphisms
AKI: acute kidney injury

## Data Availability

The following data resources are publicly available FDA Adverse Event Reporting System (FAERS) data from the Zhang et al. study [8], RNA-Seq count tables and metadata for the GTEx project from Recount3 [87], STRING database [31], and DrugBank [88]. However, the MedDRA database is a subscription database [24]. Details about data download and analysis code are discussed in “Data download and exploration” and in the “Scripts, dockers, and conda environment” Methods sections, respectively.

## Supplemental Figures

**Supplemental Figure 1:** Barplot of the top 30 female-biased adverse events based on the number of drugs with an adverse event.

**Supplemental Figure 2:** Barplot of the top 30 male-biased adverse events based on the number of drugs with an adverse event.

**Supplemental Figure 3:** Barplot of the top 30 drugs based on the number of female-biased adverse events associated with the drug.

**Supplemental Figure 4:** Barplot of the top 30 drugs based on the number of male-biased adverse events associated with the drug.

**Supplemental Figure 5:** Heatmap of the logROR of the 50 most common drugs (x-axis) and adverse events (y-axis) for both sexes. Purple logROR is female-biased and Teal logROR is male-bias.

**Supplemental Figure 6:** Bar plots of the number of drug metabolism genes in the sex-specific communities of the liver for **A)** female and **B)** male.

**Supplemental Figure 7:** We performed a one-tailed Wilcoxon test to determine if the number of core genes selected for drug metabolism was higher than for the randomly selected genes for **A)** Sex-biased liver core gene (p-value =1). **B)** male-specific liver core genes (p-value = 0.9910552) **C)** female-specific liver core genes (p-value =1).

**Supplemental Figure 8: Permutation results for the SBAE-associated drug targets’ enrichment of sex-biased expressed genes. A)** Bar plot of the female-biased sex-biased gene expression gene sets by tissue with the x-axis being the fraction of SBAE-associated drug targets with female-biased gene expression for the tissue and the number of drug targets with sex-biased gene expression. **B)** Bar plot of the male-biased sex-biased gene expression gene sets by tissue with the x-axis being the fraction of SBAE-associated drug targets with male-biased gene expression for the tissue and the number of drug targets with sex-biased expression. Permutation testing was conducted by randomly selecting either 84 drug targets genes 1,000 times with a one-tailed Wilcoxon test and with BH-multiple hypothesis test correction (ɑ = 0.05).

**Supplemental Figure 8: Permutation results for the SBAE-associated drug targets’ enrichment of sex-biased expressed genes. A)** Bar plot of the female-biased sex-biased gene expression gene sets by tissue with the x-axis being the fraction of SBAE-associated drug targets with female-biased gene expression for the tissue and the number of drug targets with sex-biased gene expression. **B)** Bar plot of the male-biased sex-biased gene expression gene sets by tissue with the x-axis being the fraction of SBAE-associated drug targets with male-biased gene expression for the tissue and the number of drug targets with sex-biased expression. Permutation testing was conducted by randomly selecting either 84 drug targets genes 1,000 times with a one-tailed Wilcoxon test and with BH-multiple hypothesis test correction (ɑ = 0.05).

**Supplemental Figure 9: Drug target *ADRA1A’s* drug-adverse event plots. A)** A bar plot of the number of male-biased drug-adverse event pairs with a drug with *ADRA1A* as drug target for each System Organ Class (SOC) adverse event term. **B)** A bar plot of the number of male-biased drug-adverse event pairs with a drug with *ADRA1A* as drug target for each Preferred Term (PT) adverse event term. **C)** A bar plot of the number of female-biased drug-adverse event pairs with a drug with *ADRA1A* as drug target for each System Organ Class (SOC) adverse event term.

**Supplemental Figure 10: Drug target *ADRA1C’s* drug-adverse event plots. A)** A bar plot of the number of male-biased drug-adverse event pairs with a drug with *ADRA1C* as drug target for each System Organ Class (SOC) adverse event term. **B)** A bar plot of the number of male-biased drug-adverse event pairs with a drug with *ADRA1C* as drug target for each Preferred Term (PT) adverse event term. **C)** A bar plot of the number of female-biased drug-adverse event pairs with a drug with *ADRA1C* as drug target for each System Organ Class (SOC) adverse event term.

**Supplemental Figure 11: Drug target *DRD1’s* drug-adverse event plots. A)** A bar plot of the number of male-biased drug-adverse event pairs with a drug with *DRD1* as drug target for each System Organ Class (SOC) adverse event term. **B)** A bar plot of the number of male-biased drug-adverse event pairs with a drug with *DRD1* as drug target for each Perferred Term (PT) adverse event term. **C)** A bar plot of the number of female-biased drug-adverse event pairs with a drug with *DRD1* as drug target for each System Organ Class (SOC) adverse event term.

**Supplemental Figure 12:** A bar plot of the number of female-biased drug-adverse event pairs with a drug with *FCGR3B* as drug target for each System Organ Class (SOC) adverse event term.

**Supplemental Figure 13: Drug target *AR’s* drug-adverse event plots. A)** A bar plot of the number of female-biased drug-adverse event pairs with a drug with *AR* as drug target for each System Organ Class (SOC) adverse event term. **B)** A bar plot of the number of male-biased drug-adverse event pairs with a drug with *AR* as drug target for each System Organ Class (SOC) adverse event term.

**Supplemental Figure 14:** A bar plot of the number of female-biased drug-adverse event pairs with a drug with *CACNA1S* as drug target for each System Organ Class (SOC) adverse event term.

**Supplemental Figure 15: Permutation results for the SBAE-associated drug targets’ enrichment of sex-specific network core genes. A)** Bar plot of the female-specific network core gene gene sets by tissue with the x-axis being the fraction of SBAE-associated drug targets with female-specific network core genes for the tissue and the number of drug targets that are female-specific network core genes. **B)** Bar plot of the male-specific network core gene gene sets by tissue with the x-axis being the fraction of SBAE-associated drug targets with male-specific network core genes for the tissue and the number of drug targets that are male-specific network core genes. Permutation testing was conducted by randomly selecting either 84 drug targets genes 1,000 times with a one-tailed Wilcoxon test and with BH-multiple hypothesis test correction (ɑ = 0.05).

## Acknowledgements

We acknowledge support from the University of Alabama at Birmingham Biological Data Science Core, RRID:SCR_021766. We would like to thank all the members of the Lasseigne Lab for their feedback on this manuscript.

## Funding

JLF and BNL were supported by R03OD030604; JLF, EFJ, and BNL were supported by R00HG009678; ADC and BNL are supported by U54OD030167; JLF, EJF, and BNL are supported by UAB funds to the Lasseigne Lab.

## Author information

### Authors and Affiliations

Department of Cell, Developmental and Integrative Biology, Heersink School of Medicine, University of Alabama at Birmingham, Birmingham, AL, 35294, USA Jennifer L. Fisher, Amanda D. Clark, Emma F. Jones, and Brittany N. Lasseigne

### Contributions

Jennifer L. Fisher: Conceptualization, Methodology, Software, Formal Analysis, Investigation, Data Curation, Writing - Original Draft, Writing - Review & Editing, Visualization. Amanda D. Clark: Validation, Writing - Review & Editing. Emma F. Jones: Validation, Writing - Review & Editing. Brittany N. Lasseigne: Conceptualization, Resources, Writing - Review & Editing, Supervision, Project administration, Funding acquisition.

### Corresponding author

Correspondence to Brittany N. Lasseigne.

## Ethics declarations

### Ethics approval and consent to participate

Not applicable.

### Consent for publication

Not applicable.

### Competing interests

The authors declare that they have no competing interests.

## Notes

https://zenodo.org/record/7938613#.ZGKFpdbMIaQ

https://zenodo.org/record/7941598#.ZGOw5tbMKJE

### Competing Interest Statement

The authors have declared no competing interest.

### Summary of Updates

We removed analyses associated with the previous version's Figure 3C and D.

