## Supplementary material for "Sex-biased gene expression and gene-regulatory networks of sex-biased adverse event drug targets and drug metabolism genes": Supp File 1

### 230510_rstudio_sex_bias_drugs_info

Jennifer Fisher

2022-12-21

```
System which operations were done on:
my laptop

GitHub Repo:
230321_JLF_Sex_bias_adverse_events

Docker:
jenfisher7/rstudio_sex_bias_drugs

Directory of operations:
~/Documents/230321_JLF_Sex_bias_adverse_events

Scripts being edited for operations:
NA

Data being used:
```

```
sessionInfo()
```

```
## R version 4.2.2 (2022-10-31)
## Platform: x86_64-pc-linux-gnu (64-bit)
## Running under: Ubuntu 22.04.1 LTS
##
## Matrix products: default
## BLAS:   /usr/lib/x86_64-linux-gnu/openblas-pthread/libblas.so.3
## LAPACK: /usr/lib/x86_64-linux-gnu/openblas-pthread/libopenblasp-r0.3.20.so
##
## locale:
##  [1] LC_CTYPE=en_US.UTF-8       LC_NUMERIC=C
##  [3] LC_TIME=en_US.UTF-8        LC_COLLATE=en_US.UTF-8
##  [5] LC_MONETARY=en_US.UTF-8    LC_MESSAGES=en_US.UTF-8
##  [7] LC_PAPER=en_US.UTF-8       LC_NAME=C
##  [9] LC_ADDRESS=C               LC_TELEPHONE=C
## [11] LC_MEASUREMENT=en_US.UTF-8 LC_IDENTIFICATION=C
##
## attached base packages:
## [1] stats     graphics  grDevices utils     datasets  methods   base
##
## loaded via a namespace (and not attached):
##  [1] digest_0.6.31   R6_2.5.1        lifecycle_1.0.3 jsonlite_1.8.4
##  [5] magrittr_2.0.3  evaluate_0.19   stringi_1.7.8   cachem_1.0.6
##  [9] rlang_1.0.6     cli_3.5.0       rstudioapi_0.14 jquerylib_0.1.4
## [13] bslib_0.4.2     vctrs_0.5.1     rmarkdown_2.19  tools_4.2.2
## [17] stringr_1.5.0   glue_1.6.2      xfun_0.36       yaml_2.3.6
## [21] fastmap_1.1.0   compiler_4.2.2  htmltools_0.5.4 knitr_1.41
## [25] sass_0.4.4
```

```
# Listing packages
installed.packages()[,c(1,3)]
```

```
##                        Package                 Version
## abind                  "abind"                 "1.4-5"
## affy                   "affy"                  "1.76.0"
## affyio                 "affyio"                "1.68.0"
## amap                   "amap"                  "0.8-19"
## annotate               "annotate"              "1.76.0"
## AnnotationDbi          "AnnotationDbi"         "1.60.0"
## AnnotationFilter       "AnnotationFilter"      "1.22.0"
## AnnotationForge        "AnnotationForge"       "1.40.0"
## AnnotationHub          "AnnotationHub"         "3.6.0"
## apcluster              "apcluster"             "1.4.10"
## ape                    "ape"                   "5.6-2"
## aplot                  "aplot"                 "0.1.9"
## ashr                   "ashr"                  "2.2-54"
## askpass                "askpass"               "1.1"
## assertthat             "assertthat"            "0.2.1"
## backports              "backports"             "1.4.1"
## base64                 "base64"                "2.0.1"
## base64enc              "base64enc"             "0.1-3"
## base64url              "base64url"             "1.4"
## bayestestR             "bayestestR"            "0.13.0"
## beanplot               "beanplot"              "1.3.1"
## bench                  "bench"                 "1.1.2"
## BgeeDB                 "BgeeDB"                "2.24.0"
## BH                     "BH"                    "1.78.0-0"
## Biobase                "Biobase"               "2.58.0"
## BiocBaseUtils          "BiocBaseUtils"         "1.0.0"
## BiocFileCache          "BiocFileCache"         "2.6.0"
## BiocGenerics           "BiocGenerics"          "0.44.0"
## BiocIO                 "BiocIO"                "1.8.0"
## BiocManager            "BiocManager"           "1.30.19"
## BiocParallel           "BiocParallel"          "1.32.5"
## BiocVersion            "BiocVersion"           "3.16.0"
## biomaRt                "biomaRt"               "2.54.0"
## Biostrings             "Biostrings"            "2.66.0"
## bit                    "bit"                   "4.0.5"
## bit64                  "bit64"                 "4.0.5"
## bitops                 "bitops"                "1.0-7"
## biwt                   "biwt"                  "1.0.1"
## blob                   "blob"                  "1.2.3"
## brew                   "brew"                  "1.0-8"
## brio                   "brio"                  "1.1.3"
## broom                  "broom"                 "1.0.2"
## broom.mixed            "broom.mixed"           "0.2.9.4"
## bslib                  "bslib"                 "0.4.2"
## bumphunter             "bumphunter"            "1.40.0"
## BumpyMatrix            "BumpyMatrix"           "1.6.0"
## cachem                 "cachem"                "1.0.6"
## callr                  "callr"                 "3.7.3"
## car                    "car"                   "3.1-1"
## carData                "carData"               "3.0-5"
## caret                  "caret"                 "6.0-93"
```

```
## Category                "Category"                "2.64.0"
## caTools                 "caTools"                 "1.18.2"
## celestial               "celestial"               "1.4.6"
## cellranger              "cellranger"              "1.1.0"
## checkmate               "checkmate"               "2.1.0"
## ChemmineR               "ChemmineR"               "3.50.0"
## chron                   "chron"                   "2.3-58"
## circlize                "circlize"                "0.4.15"
## classInt                "classInt"                "0.4-8"
## cli                     "cli"                     "3.5.0"
## clipr                   "clipr"                   "0.8.0"
## clock                   "clock"                   "0.6.1"
## clue                    "clue"                    "0.3-63"
## clusterProfiler         "clusterProfiler"         "4.6.0"
## coda                    "coda"                    "0.19-4"
## CoGAPS                  "CoGAPS"                  "3.18.0"
## cogena                  "cogena"                  "1.32.0"
## colorspace              "colorspace"              "2.0-3"
## combinat                "combinat"                "0.0-8"
## commonmark              "commonmark"              "1.8.1"
## ComplexHeatmap          "ComplexHeatmap"          "2.14.0"
## ComplexUpset            "ComplexUpset"            "1.3.3"
## conflicted              "conflicted"              "1.1.0"
## coop                    "coop"                    "0.6-3"
## CoreGx                  "CoreGx"                  "2.2.0"
## corrplot                "corrplot"                "0.92"
## cowplot                 "cowplot"                 "1.1.1"
## cpp11                   "cpp11"                   "0.4.3"
## crayon                  "crayon"                  "1.5.2"
## credentials             "credentials"             "1.3.2"
## crosstalk               "crosstalk"               "1.2.0"
## curl                    "curl"                    "4.3.3"
## customCMPdb             "customCMPdb"             "1.8.0"
## data.table              "data.table"              "1.14.6"
## datawizard              "datawizard"              "0.6.5"
## DBI                     "DBI"                     "1.1.3"
## dbplyr                  "dbplyr"                  "2.2.1"
## DelayedArray            "DelayedArray"            "0.24.0"
## DelayedMatrixStats      "DelayedMatrixStats"      "1.20.0"
## dendextend              "dendextend"              "1.16.0"
## DEoptimR                "DEoptimR"                "1.0-11"
## desc                    "desc"                    "1.4.2"
## DESeq2                  "DESeq2"                  "1.38.2"
## devtools                "devtools"                "2.4.5"
## DEXSeq                  "DEXSeq"                  "1.44.0"
## dials                   "dials"                   "1.1.0"
## DiceDesign              "DiceDesign"              "1.9"
## diffobj                 "diffobj"                 "0.3.5"
## digest                  "digest"                  "0.6.31"
## discrim                 "discrim"                 "1.0.0"
## docopt                  "docopt"                  "0.7.1"
## doParallel              "doParallel"              "1.0.17"
```

```
## doRNG                 "doRNG"                 "1.8.3"
## DOSE                  "DOSE"                  "3.24.2"
## dotwhisker            "dotwhisker"            "0.7.4"
## downlit               "downlit"               "0.4.2"
## downloader            "downloader"            "0.4"
## dplyr                 "dplyr"                 "1.0.10"
## drugbankR             "drugbankR"             "1.5"
## DT                    "DT"                    "0.26"
## dtplyr                "dtplyr"                "1.2.2"
## e1071                 "e1071"                 "1.7-12"
## earth                 "earth"                 "5.3.1"
## edgeR                 "edgeR"                 "3.40.1"
## ellipse               "ellipse"               "0.4.3"
## ellipsis              "ellipsis"              "0.3.2"
## emmeans               "emmeans"               "1.8.3"
## enrichplot            "enrichplot"            "1.18.3"
## EnsDb.Hsapiens.v75    "EnsDb.Hsapiens.v75"    "2.99.0"
## ensembldb             "ensembldb"             "2.22.0"
## estimability          "estimability"          "1.4.1"
## etrunct               "etrunct"               "0.1"
## evaluate              "evaluate"              "0.19"
## ExperimentHub         "ExperimentHub"         "2.6.0"
## factoextra            "factoextra"            "1.0.7"
## FactoMineR            "FactoMineR"            "2.7"
## fansi                 "fansi"                 "1.0.3"
## farver                "farver"                "2.1.1"
## fastcluster           "fastcluster"           "1.2.3"
## fastmap               "fastmap"               "1.1.0"
## fastmatch             "fastmatch"             "1.1-3"
## fgsea                 "fgsea"                 "1.24.0"
## filelock              "filelock"              "1.0.2"
## flashClust            "flashClust"            "1.01-2"
## fmcsR                 "fmcsR"                 "1.40.0"
## fontawesome           "fontawesome"           "0.4.0"
## forcats               "forcats"               "0.5.2"
## foreach               "foreach"               "1.5.2"
## formatR               "formatR"               "1.13"
## Formula               "Formula"               "1.2-4"
## fs                    "fs"                    "1.5.2"
## furrr                 "furrr"                 "0.3.1"
## futile.logger         "futile.logger"         "1.4.3"
## futile.options        "futile.options"        "1.0.1"
## future                "future"                "1.30.0"
## future.apply          "future.apply"          "1.10.0"
## gargle                "gargle"                "1.2.1"
## gbm                   "gbm"                   "2.1.8.1"
## genefilter            "genefilter"            "1.80.2"
## geneplotter           "geneplotter"           "1.76.0"
## generics              "generics"              "0.1.3"
## GenomeInfoDb          "GenomeInfoDb"          "1.34.4"
## GenomeInfoDbData      "GenomeInfoDbData"      "1.2.9"
## GenomicAlignments     "GenomicAlignments"     "1.34.0"
```

```
## GenomicFeatures         "GenomicFeatures"         "1.50.3"
## GenomicRanges           "GenomicRanges"           "1.50.2"
## GEOquery                "GEOquery"                "2.66.0"
## gert                    "gert"                    "1.9.1"
## GetoptLong              "GetoptLong"              "1.0.5"
## ggalluvial              "ggalluvial"              "0.12.3"
## ggdendro                "ggdendro"                "0.1.23"
## ggforce                 "ggforce"                 "0.4.1"
## ggfun                   "ggfun"                   "0.0.9"
## ggnewscale              "ggnewscale"              "0.4.8"
## ggplot2                 "ggplot2"                 "3.4.0"
## ggplotify               "ggplotify"               "0.1.0"
## ggpubr                  "ggpubr"                  "0.5.0"
## ggraph                  "ggraph"                  "2.1.0"
## ggrepel                 "ggrepel"                 "0.9.2"
## ggsci                   "ggsci"                   "2.9"
## ggsignif                "ggsignif"                "0.6.4"
## ggstance                "ggstance"                "0.3.6"
## ggtree                  "ggtree"                  "3.6.2"
## gh                      "gh"                      "1.3.1"
## gitcreds                "gitcreds"                "0.1.2"
## glmnet                  "glmnet"                  "4.1-6"
## GlobalOptions           "GlobalOptions"           "0.1.2"
## globals                 "globals"                 "0.16.2"
## glue                    "glue"                    "1.6.2"
## GO.db                   "GO.db"                   "3.16.0"
## googledrive             "googledrive"             "2.0.0"
## googlesheets4           "googlesheets4"           "1.0.1"
## GOSemSim                "GOSemSim"                "2.24.0"
## GOstats                 "GOstats"                 "2.64.0"
## gower                   "gower"                   "1.0.1"
## GPfit                   "GPfit"                   "1.0-8"
## gplots                  "gplots"                  "3.1.3"
## gprofiler2              "gprofiler2"              "0.2.1"
## graph                   "graph"                   "1.76.0"
## graphlayouts            "graphlayouts"            "0.8.4"
## gridBase                "gridBase"                "0.4-7"
## gridExtra               "gridExtra"               "2.3"
## gridGraphics            "gridGraphics"            "0.5-1"
## GSEABase                "GSEABase"                "1.60.0"
## gson                    "gson"                    "0.0.9"
## gsubfn                  "gsubfn"                  "0.7"
## gtable                  "gtable"                  "0.3.1"
## gtools                  "gtools"                  "3.9.4"
## hardhat                 "hardhat"                 "1.2.0"
## hash                    "hash"                    "2.2.6.2"
## haven                   "haven"                   "2.5.1"
## HDF5Array               "HDF5Array"               "1.26.0"
## HDO.db                  "HDO.db"                  "0.99.1"
## here                    "here"                    "1.0.1"
## hexbin                  "hexbin"                  "1.28.2"
## highr                   "highr"                   "0.10"
```

```
## hms                    "hms"                    "1.1.2"
## htmltools              "htmltools"              "0.5.4"
## htmlwidgets            "htmlwidgets"            "1.6.0"
## httpuv                 "httpuv"                 "1.6.7"
## httr                   "httr"                   "1.4.4"
## hwriter                "hwriter"                "1.3.2.1"
## ids                    "ids"                    "1.0.1"
## igraph                 "igraph"                 "1.3.5"
## illuminaio             "illuminaio"             "0.40.0"
## infer                  "infer"                  "1.0.4"
## ini                    "ini"                    "0.3.1"
## insight                "insight"                "0.18.8"
## interactiveDisplayBase "interactiveDisplayBase" "1.36.0"
## inum                   "inum"                   "1.0-4"
## invgamma               "invgamma"               "1.1"
## ipred                  "ipred"                  "0.9-13"
## IRanges                "IRanges"                "2.32.0"
## IRdisplay              "IRdisplay"              "1.1"
## IRkernel               "IRkernel"               "1.3.1"
## irlba                  "irlba"                  "2.3.5.1"
## isoband                "isoband"                "0.2.7"
## iterators              "iterators"              "1.0.14"
## jquerylib              "jquerylib"              "0.1.4"
## jsonlite               "jsonlite"               "1.8.4"
## KEGGREST               "KEGGREST"               "1.38.0"
## kernlab                "kernlab"                "0.9-31"
## klaR                   "klaR"                   "1.7-1"
## knitr                  "knitr"                  "1.41"
## kohonen                "kohonen"                "3.0.11"
## labeling               "labeling"               "0.4.2"
## labelled               "labelled"               "2.10.0"
## lambda.r               "lambda.r"               "1.2.4"
## later                  "later"                  "1.3.0"
## lava                   "lava"                   "1.7.0"
## lazyeval               "lazyeval"               "0.2.2"
## leaps                  "leaps"                  "3.1"
## lhs                    "lhs"                    "1.1.6"
## libcoin                "libcoin"                "1.0-9"
## LiblineaR              "LiblineaR"              "2.10-22"
## lifecycle              "lifecycle"              "1.0.3"
## limma                  "limma"                  "3.54.0"
## listenv                "listenv"                "0.9.0"
## littler                "littler"                "0.3.17"
## lme4                   "lme4"                   "1.1-31"
## locfit                 "locfit"                 "1.5-9.7"
## lsa                    "lsa"                    "0.73.3"
## lubridate              "lubridate"              "1.9.0"
## magicaxis              "magicaxis"              "2.2.14"
## magrittr               "magrittr"               "2.0.3"
## mapproj                "mapproj"                "1.2.9"
## maps                   "maps"                   "3.4.1"
## margins                "margins"                "0.3.26"
```

```
## markdown               "markdown"               "1.4"
## marray                 "marray"                 "1.76.0"
## mashr                  "mashr"                  "0.2.69"
## MASS                   "MASS"                   "7.3-58.1"
## matrixcalc             "matrixcalc"             "1.0-6"
## MatrixGenerics         "MatrixGenerics"         "1.10.0"
## MatrixModels           "MatrixModels"           "0.5-1"
## matrixStats            "matrixStats"            "0.63.0"
## mclust                 "mclust"                 "6.0.0"
## memoise                "memoise"                "2.0.1"
## mime                   "mime"                   "0.12"
## minfi                  "minfi"                  "1.44.0"
## miniUI                 "miniUI"                 "0.1.1.1"
## minqa                  "minqa"                  "1.2.5"
## mixsqp                 "mixsqp"                 "0.3-48"
## modeldata              "modeldata"              "1.0.1"
## modelenv               "modelenv"               "0.1.0"
## ModelMetrics           "ModelMetrics"           "1.2.2.2"
## modelr                 "modelr"                 "0.1.10"
## multcompView           "multcompView"           "0.1-8"
## MultiAssayExperiment   "MultiAssayExperiment"   "1.24.0"
## multtest               "multtest"               "2.54.0"
## munsell                "munsell"                "0.5.0"
## mvtnorm                "mvtnorm"                "1.1-3"
## naivebayes             "naivebayes"             "0.9.7"
## netZooR                "netZooR"                "1.2.1"
## NISTunits              "NISTunits"              "1.0.1"
## nloptr                 "nloptr"                 "2.0.3"
## NMF                    "NMF"                    "0.25"
## nnet                   "nnet"                   "7.3-18"
## nor1mix                "nor1mix"                "1.3-0"
## numDeriv               "numDeriv"               "2016.8-1.1"
## openssl                "openssl"                "2.0.5"
## org.Hs.eg.db           "org.Hs.eg.db"           "3.16.0"
## pamr                   "pamr"                   "1.56.1"
## pandaR                 "pandaR"                 "1.30.0"
## parallelly             "parallelly"             "1.33.0"
## parameters             "parameters"             "0.20.0"
## parsnip                "parsnip"                "1.0.3"
## partykit               "partykit"               "1.2-16"
## pasilla                "pasilla"                "1.26.0"
## patchwork              "patchwork"              "1.1.2"
## pbdZMQ                 "pbdZMQ"                 "0.3-8"
## pbkrtest               "pbkrtest"               "0.5.1"
## penalized              "penalized"              "0.9-52"
## permute                "permute"                "0.9-7"
## PharmacoGx             "PharmacoGx"             "3.2.0"
## pheatmap               "pheatmap"               "1.0.12"
## piano                  "piano"                  "2.14.0"
## pillar                 "pillar"                 "1.8.1"
## pkgbuild               "pkgbuild"               "1.3.1"
## pkgconfig              "pkgconfig"              "2.0.3"
```

```
## pkgdown             "pkgdown"             "2.0.6"
## pkgload             "pkgload"             "1.3.1"
## PLIER               "PLIER"               "0.99.0"
## plogr               "plogr"               "0.2.0"
## plotly              "plotly"              "4.10.1"
## plotmo              "plotmo"              "3.6.2"
## plotrix             "plotrix"             "3.8-2"
## plyr                "plyr"                "1.8.8"
## png                 "png"                 "0.1-8"
## polyclip            "polyclip"            "1.10-4"
## polynom             "polynom"             "1.4-1"
## pracma              "pracma"              "2.4.2"
## praise              "praise"              "1.0.0"
## prediction          "prediction"          "0.3.14"
## preprocessCore      "preprocessCore"      "1.60.1"
## prettyunits         "prettyunits"         "1.1.1"
## pROC                "pROC"                "1.18.0"
## processx            "processx"            "3.8.0"
## prodlim             "prodlim"             "2019.11.13"
## profmem             "profmem"             "0.6.0"
## profvis             "profvis"             "0.3.7"
## progress            "progress"            "1.2.2"
## progressr           "progressr"           "0.12.0"
## projectR            "projectR"            "1.14.0"
## ProliferativeIndex  "ProliferativeIndex"  "1.0.1"
## promises            "promises"            "1.2.0.1"
## ProtGenerics        "ProtGenerics"        "1.30.0"
## proto               "proto"               "1.0.0"
## proxy               "proxy"               "0.4-27"
## ps                  "ps"                  "1.7.2"
## purrr               "purrr"               "1.0.0"
## quadprog            "quadprog"            "1.5-8"
## quantreg            "quantreg"            "5.94"
## quantro             "quantro"             "1.32.0"
## questionr           "questionr"           "0.7.7"
## qvalue              "qvalue"              "2.30.0"
## R.cache             "R.cache"             "0.16.0"
## R.methodsS3         "R.methodsS3"         "1.8.2"
## R.oo                "R.oo"                "1.25.0"
## R.utils             "R.utils"             "2.12.2"
## R6                  "R6"                  "2.5.1"
## ragg                "ragg"                "1.2.4"
## randomForest        "randomForest"        "4.7-1.1"
## ranger              "ranger"              "0.14.1"
## RANN                "RANN"                "2.6.1"
## rappdirs            "rappdirs"            "0.3.3"
## RBGL                "RBGL"                "1.74.0"
## rcmdcheck           "rcmdcheck"           "1.4.0"
## RColorBrewer        "RColorBrewer"        "1.1-3"
## Rcpp                "Rcpp"                "1.0.9"
## RcppArmadillo       "RcppArmadillo"       "0.11.4.2.1"
## RcppEigen           "RcppEigen"           "0.3.3.9.3"
```

```
## RcppGSL             "RcppGSL"             "0.3.12"
## RcppTOML            "RcppTOML"            "0.1.7"
## RCurl               "RCurl"               "1.98-1.9"
## RCy3                "RCy3"                "2.18.0"
## reactome.db         "reactome.db"         "1.82.0"
## readr               "readr"               "2.1.3"
## readxl              "readxl"              "1.4.1"
## recipes             "recipes"             "1.0.3"
## recount3            "recount3"            "1.8.0"
## registry            "registry"            "0.5-1"
## relations           "relations"           "0.6-12"
## rematch             "rematch"             "1.0.1"
## rematch2            "rematch2"            "2.1.2"
## remotes             "remotes"             "2.4.2"
## repr                "repr"                "1.1.4"
## reprex              "reprex"              "2.0.2"
## reshape             "reshape"             "0.8.9"
## reshape2            "reshape2"            "1.4.4"
## restfulr            "restfulr"            "0.0.15"
## reticulate          "reticulate"          "1.26"
## Rgraphviz           "Rgraphviz"           "2.42.0"
## rhdf5               "rhdf5"               "2.42.0"
## rhdf5filters        "rhdf5filters"        "1.10.0"
## Rhdf5lib            "Rhdf5lib"            "1.20.0"
## Rhtslib             "Rhtslib"             "2.0.0"
## rJava               "rJava"               "1.0-6"
## rjson               "rjson"               "0.2.21"
## RJSONIO             "RJSONIO"             "1.3-1.6"
## rlang               "rlang"               "1.0.6"
## rmarkdown           "rmarkdown"           "2.19"
## rmeta               "rmeta"               "3.0"
## rngtools            "rngtools"            "1.5.2"
## robustbase          "robustbase"          "0.95-0"
## ROCR                "ROCR"                "1.0-11"
## roxygen2            "roxygen2"            "7.2.1"
## rpart               "rpart"               "4.1.19"
## rprojroot           "rprojroot"           "2.0.3"
## rsample             "rsample"             "1.1.1"
## Rsamtools           "Rsamtools"           "2.14.0"
## RSQLite             "RSQLite"             "2.2.20"
## rstatix             "rstatix"             "0.7.1"
## rstudioapi          "rstudioapi"          "0.14"
## rsvd                "rsvd"                "1.0.5"
## rsvg                "rsvg"                "2.4.0"
## rtracklayer         "rtracklayer"         "1.58.0"
## RUnit               "RUnit"               "0.4.32"
## rversions           "rversions"           "2.1.2"
## rvest               "rvest"               "1.0.3"
## RWeka               "RWeka"               "0.4-44"
## RWekajars           "RWekajars"           "3.9.3-2"
## S4Vectors           "S4Vectors"           "0.36.1"
## sass                "sass"                "0.4.4"
```

```
## scales                   "scales"                   "1.2.1"
## scatterpie               "scatterpie"               "0.1.8"
## scatterplot3d            "scatterplot3d"            "0.3-42"
## scrime                   "scrime"                   "1.3.5"
## selectr                  "selectr"                  "0.4-2"
## sessioninfo              "sessioninfo"              "1.2.2"
## sets                     "sets"                     "1.0-21"
## shadowtext               "shadowtext"               "0.1.2"
## shape                    "shape"                    "1.4.6"
## shiny                    "shiny"                    "1.7.4"
## shinydashboard           "shinydashboard"           "0.7.2"
## shinyjs                  "shinyjs"                  "2.1.0"
## siggenes                 "siggenes"                 "1.72.0"
## signatureSearch          "signatureSearch"          "1.11.1"
## signatureSearchData      "signatureSearchData"      "1.12.0"
## SingleCellExperiment     "SingleCellExperiment"     "1.20.0"
## skimr                    "skimr"                    "2.1.5"
## slam                     "slam"                     "0.1-50"
## slider                   "slider"                   "0.3.0"
## sm                       "sm"                       "2.2-5.7.1"
## snow                     "snow"                     "0.4-4"
## SnowballC                "SnowballC"                "0.7.0"
## softImpute               "softImpute"               "1.4-1"
## sourcetools              "sourcetools"              "0.1.7"
## SparseM                  "SparseM"                  "1.81"
## sparseMatrixStats        "sparseMatrixStats"        "1.10.0"
## sqldf                    "sqldf"                    "0.4-11"
## SQUAREM                  "SQUAREM"                  "2021.1"
## statmod                  "statmod"                  "1.4.37"
## STRINGdb                 "STRINGdb"                 "2.10.0"
## stringi                  "stringi"                  "1.7.8"
## stringr                  "stringr"                  "1.5.0"
## styler                   "styler"                   "1.8.1"
## SummarizedExperiment     "SummarizedExperiment"     "1.28.0"
## sys                      "sys"                      "3.4.1"
## systemfonts              "systemfonts"              "1.0.4"
## TeachingDemos            "TeachingDemos"            "2.12"
## testthat                 "testthat"                 "3.1.5"
## textshaping              "textshaping"              "0.3.6"
## TFEA.ChIP                "TFEA.ChIP"                "1.18.0"
## tibble                   "tibble"                   "3.1.8"
## tidygraph                "tidygraph"                "1.2.2"
## tidymodels               "tidymodels"               "1.0.0"
## tidyr                    "tidyr"                    "1.2.1"
## tidyselect               "tidyselect"               "1.2.0"
## tidytree                 "tidytree"                 "0.4.2"
## tidyverse                "tidyverse"                "1.3.2"
## timechange               "timechange"               "0.1.1"
## timeDate                 "timeDate"                 "4021.107"
## tinytex                  "tinytex"                  "0.43"
## topGO                    "topGO"                    "2.50.0"
## treeio                   "treeio"                   "1.22.0"
```

```
## truncnorm          "truncnorm"          "1.0-8"
## tune               "tune"               "1.0.1"
## tweenr             "tweenr"             "2.0.2"
## tzdb               "tzdb"               "0.3.0"
## uchardet           "uchardet"           "1.1.1"
## urlchecker         "urlchecker"         "1.0.1"
## usethis            "usethis"            "2.1.6"
## utf8               "utf8"               "1.2.2"
## uuid               "uuid"               "1.1-0"
## vctrs              "vctrs"              "0.5.1"
## vegan              "vegan"              "2.6-4"
## VennDiagram        "VennDiagram"        "1.7.3"
## viridis            "viridis"            "0.6.2"
## viridisLite        "viridisLite"        "0.4.1"
## visNetwork         "visNetwork"         "2.1.2"
## vroom              "vroom"              "1.6.0"
## waldo              "waldo"              "0.4.0"
## warp               "warp"               "0.2.0"
## whisker            "whisker"            "0.4"
## withr              "withr"              "2.5.0"
## workflows          "workflows"          "1.1.2"
## workflowsets       "workflowsets"       "1.0.0"
## xfun               "xfun"               "0.36"
## xgboost            "xgboost"            "1.6.0.1"
## XML                "XML"                "3.99-0.13"
## xml2               "xml2"               "1.3.3"
## xopen              "xopen"              "1.0.0"
## xtable             "xtable"             "1.8-4"
## XVector            "XVector"            "0.38.0"
## yaml               "yaml"               "2.3.6"
## yardstick          "yardstick"          "1.1.0"
## yarn               "yarn"               "1.24.0"
## yulab.utils        "yulab.utils"        "0.0.6"
## zip                "zip"                "2.2.2"
## zlibbioc           "zlibbioc"           "1.44.0"
## base               "base"               "4.2.2"
## boot               "boot"               "1.3-28"
## class              "class"              "7.3-20"
## cluster            "cluster"            "2.1.4"
## codetools          "codetools"          "0.2-18"
## compiler           "compiler"           "4.2.2"
## datasets           "datasets"           "4.2.2"
## foreign            "foreign"            "0.8-83"
## graphics           "graphics"           "4.2.2"
## grDevices          "grDevices"          "4.2.2"
## grid               "grid"               "4.2.2"
## KernSmooth         "KernSmooth"         "2.23-20"
## lattice            "lattice"            "0.20-45"
## MASS               "MASS"               "7.3-58.1"
## Matrix             "Matrix"             "1.5-1"
## methods            "methods"            "4.2.2"
## mgcv               "mgcv"               "1.8-41"
```

```
## nlme                        "nlme"                    "3.1-160"
## nnet                        "nnet"                    "7.3-18"
## parallel                    "parallel"                "4.2.2"
## rpart                       "rpart"                   "4.1.19"
## spatial                     "spatial"                 "7.3-15"
## splines                     "splines"                 "4.2.2"
## stats                       "stats"                   "4.2.2"
## stats4                      "stats4"                  "4.2.2"
## survival                    "survival"                "3.4-0"
## tcltk                       "tcltk"                   "4.2.2"
## tools                       "tools"                   "4.2.2"
## utils                       "utils"                   "4.2.2"
```

### 230510_rstudio_sex_bias_drugs_singularity

Jennifer Fisher

### 05/10/2023

```
System which operations were done on:
cheaha

GitHub Repo:
230321_JLF_Sex_bias_adverse_events

Directory of operations:
/data/project/lasseigne_lab/JLF_scratch/230321_JLF_Sex_bias_adverse_events

GitHub Repo:
230321_JLF_Sex_bias_adverse_events

Docker:
rstudio_sex_bias_drugs (singularity)
```

```
sessionInfo()
```

```
## R version 4.2.2 (2022-10-31)
## Platform: x86_64-pc-linux-gnu (64-bit)
## Running under: Ubuntu 22.04.1 LTS
##
## Matrix products: default
## BLAS:   /usr/lib/x86_64-linux-gnu/openblas-pthread/libblas.so.3
## LAPACK: /usr/lib/x86_64-linux-gnu/openblas-pthread/libopenblasp-r0.3.20.so
##
## locale:
##  [1] LC_CTYPE=en_US.UTF-8       LC_NUMERIC=C
##  [3] LC_TIME=en_US.UTF-8        LC_COLLATE=en_US.UTF-8
##  [5] LC_MONETARY=en_US.UTF-8    LC_MESSAGES=en_US.UTF-8
##  [7] LC_PAPER=en_US.UTF-8       LC_NAME=C
##  [9] LC_ADDRESS=C               LC_TELEPHONE=C
## [11] LC_MEASUREMENT=en_US.UTF-8 LC_IDENTIFICATION=C
##
## attached base packages:
## [1] stats     graphics  grDevices utils     datasets  methods   base
##
## loaded via a namespace (and not attached):
##  [1] digest_0.6.31   R6_2.5.1        lifecycle_1.0.3 jsonlite_1.8.4
##  [5] magrittr_2.0.3  evaluate_0.19   stringi_1.7.8   cachem_1.0.6
##  [9] rlang_1.0.6     cli_3.5.0       rstudioapi_0.14 jquerylib_0.1.4
## [13] bslib_0.4.2     vctrs_0.5.1     rmarkdown_2.19  tools_4.2.2
## [17] stringr_1.5.0   glue_1.6.2      xfun_0.36       yaml_2.3.6
## [21] fastmap_1.1.0   compiler_4.2.2  htmltools_0.5.4 knitr_1.41
## [25] sass_0.4.4
```

```
# Listing packages
installed.packages()[,c(1,3)]
```

```
##                         Package                   Version
## abind                   "abind"                   "1.4-5"
## affy                    "affy"                    "1.76.0"
## affyio                  "affyio"                  "1.68.0"
## amap                    "amap"                    "0.8-19"
## annotate                "annotate"                "1.76.0"
## AnnotationDbi           "AnnotationDbi"           "1.60.0"
## AnnotationFilter        "AnnotationFilter"        "1.22.0"
## AnnotationForge         "AnnotationForge"         "1.40.0"
## AnnotationHub           "AnnotationHub"           "3.6.0"
## apcluster               "apcluster"               "1.4.10"
## ape                     "ape"                     "5.6-2"
## aplot                   "aplot"                   "0.1.9"
## ashr                    "ashr"                    "2.2-54"
## askpass                 "askpass"                 "1.1"
## assertthat              "assertthat"              "0.2.1"
## backports               "backports"               "1.4.1"
## base64                  "base64"                  "2.0.1"
## base64enc               "base64enc"               "0.1-3"
## base64url               "base64url"               "1.4"
## bayestestR              "bayestestR"              "0.13.0"
## beanplot                "beanplot"                "1.3.1"
## bench                   "bench"                   "1.1.2"
## BgeeDB                  "BgeeDB"                  "2.24.0"
## BH                      "BH"                      "1.78.0-0"
## Biobase                 "Biobase"                 "2.58.0"
## BiocBaseUtils           "BiocBaseUtils"           "1.0.0"
## BiocFileCache           "BiocFileCache"           "2.6.0"
## BiocGenerics            "BiocGenerics"            "0.44.0"
## BiocIO                  "BiocIO"                  "1.8.0"
## BiocManager             "BiocManager"             "1.30.19"
## BiocParallel            "BiocParallel"            "1.32.5"
## BiocVersion             "BiocVersion"             "3.16.0"
## biomaRt                 "biomaRt"                 "2.54.0"
## Biostrings              "Biostrings"              "2.66.0"
## bit                     "bit"                     "4.0.5"
## bit64                   "bit64"                   "4.0.5"
## bitops                  "bitops"                  "1.0-7"
## biwt                    "biwt"                    "1.0.1"
## blob                    "blob"                    "1.2.3"
## brew                    "brew"                    "1.0-8"
## brio                    "brio"                    "1.1.3"
## broom                   "broom"                   "1.0.2"
## broom.mixed             "broom.mixed"             "0.2.9.4"
## bslib                   "bslib"                   "0.4.2"
## bumphunter              "bumphunter"              "1.40.0"
## BumpyMatrix             "BumpyMatrix"             "1.6.0"
## cachem                  "cachem"                  "1.0.6"
## callr                   "callr"                   "3.7.3"
## car                     "car"                     "3.1-1"
## carData                 "carData"                 "3.0-5"
## caret                   "caret"                   "6.0-93"
## Category                "Category"                "2.64.0"
## caTools                 "caTools"                 "1.18.2"
```

```
## celestial            "celestial"           "1.4.6"
## cellranger           "cellranger"          "1.1.0"
## checkmate            "checkmate"           "2.1.0"
## ChemmineR            "ChemmineR"           "3.50.0"
## chron                "chron"               "2.3-58"
## circlize             "circlize"            "0.4.15"
## classInt             "classInt"            "0.4-8"
## cli                  "cli"                 "3.5.0"
## clipr                "clipr"               "0.8.0"
## clock                "clock"               "0.6.1"
## clue                 "clue"                "0.3-63"
## clusterProfiler      "clusterProfiler"     "4.6.0"
## coda                 "coda"                "0.19-4"
## CoGAPS               "CoGAPS"              "3.18.0"
## cogena               "cogena"              "1.32.0"
## colorspace           "colorspace"          "2.0-3"
## combinat             "combinat"            "0.0-8"
## commonmark           "commonmark"          "1.8.1"
## ComplexHeatmap       "ComplexHeatmap"      "2.14.0"
## ComplexUpset         "ComplexUpset"        "1.3.3"
## conflicted           "conflicted"          "1.1.0"
## coop                 "coop"                "0.6-3"
## CoreGx               "CoreGx"              "2.2.0"
## corrplot             "corrplot"            "0.92"
## cowplot              "cowplot"             "1.1.1"
## cpp11                "cpp11"               "0.4.3"
## crayon               "crayon"              "1.5.2"
## credentials          "credentials"         "1.3.2"
## crosstalk            "crosstalk"           "1.2.0"
## curl                 "curl"                "4.3.3"
## customCMPdb          "customCMPdb"         "1.8.0"
## data.table           "data.table"          "1.14.6"
## datawizard           "datawizard"          "0.6.5"
## DBI                  "DBI"                 "1.1.3"
## dbplyr               "dbplyr"              "2.2.1"
## DelayedArray         "DelayedArray"        "0.24.0"
## DelayedMatrixStats   "DelayedMatrixStats"  "1.20.0"
## dendextend           "dendextend"          "1.16.0"
## DEoptimR             "DEoptimR"            "1.0-11"
## desc                 "desc"                "1.4.2"
## DESeq2               "DESeq2"              "1.38.2"
## devtools             "devtools"            "2.4.5"
## DEXSeq               "DEXSeq"              "1.44.0"
## dials                "dials"               "1.1.0"
## DiceDesign           "DiceDesign"          "1.9"
## diffobj              "diffobj"             "0.3.5"
## digest               "digest"              "0.6.31"
## discrim              "discrim"             "1.0.0"
## docopt               "docopt"              "0.7.1"
## doParallel           "doParallel"          "1.0.17"
## doRNG                "doRNG"               "1.8.3"
## DOSE                 "DOSE"                "3.24.2"
## dotwhisker           "dotwhisker"          "0.7.4"
## downlit              "downlit"             "0.4.2"
## downloader           "downloader"          "0.4"
```

```
## dplyr                "dplyr"                "1.0.10"
## drugbankR            "drugbankR"            "1.5"
## DT                   "DT"                   "0.26"
## dtplyr               "dtplyr"               "1.2.2"
## e1071                "e1071"                "1.7-12"
## earth                "earth"                "5.3.1"
## edgeR                "edgeR"                "3.40.1"
## ellipse              "ellipse"              "0.4.3"
## ellipsis             "ellipsis"             "0.3.2"
## emmeans              "emmeans"              "1.8.3"
## enrichplot           "enrichplot"           "1.18.3"
## EnsDb.Hsapiens.v75   "EnsDb.Hsapiens.v75"   "2.99.0"
## ensembldb            "ensembldb"            "2.22.0"
## estimability         "estimability"         "1.4.1"
## etrunct              "etrunct"              "0.1"
## evaluate             "evaluate"             "0.19"
## ExperimentHub        "ExperimentHub"        "2.6.0"
## factoextra           "factoextra"           "1.0.7"
## FactoMineR           "FactoMineR"           "2.7"
## fansi                "fansi"                "1.0.3"
## farver               "farver"               "2.1.1"
## fastcluster          "fastcluster"          "1.2.3"
## fastmap              "fastmap"              "1.1.0"
## fastmatch            "fastmatch"            "1.1-3"
## fgsea                "fgsea"                "1.24.0"
## filelock             "filelock"             "1.0.2"
## flashClust           "flashClust"           "1.01-2"
## fmcsR                "fmcsR"                "1.40.0"
## fontawesome          "fontawesome"          "0.4.0"
## forcats              "forcats"              "0.5.2"
## foreach              "foreach"              "1.5.2"
## formatR              "formatR"              "1.13"
## Formula              "Formula"              "1.2-4"
## fs                   "fs"                   "1.5.2"
## furrr                "furrr"                "0.3.1"
## futile.logger        "futile.logger"        "1.4.3"
## futile.options       "futile.options"       "1.0.1"
## future               "future"               "1.30.0"
## future.apply         "future.apply"         "1.10.0"
## gargle               "gargle"               "1.2.1"
## gbm                  "gbm"                  "2.1.8.1"
## genefilter           "genefilter"           "1.80.2"
## geneplotter          "geneplotter"          "1.76.0"
## generics             "generics"             "0.1.3"
## GenomeInfoDb         "GenomeInfoDb"         "1.34.4"
## GenomeInfoDbData     "GenomeInfoDbData"     "1.2.9"
## GenomicAlignments    "GenomicAlignments"    "1.34.0"
## GenomicFeatures      "GenomicFeatures"      "1.50.3"
## GenomicRanges        "GenomicRanges"        "1.50.2"
## GEOquery             "GEOquery"             "2.66.0"
## gert                 "gert"                 "1.9.1"
## GetoptLong           "GetoptLong"           "1.0.5"
## ggalluvial           "ggalluvial"           "0.12.3"
## ggdendro             "ggdendro"             "0.1.23"
## ggforce              "ggforce"              "0.4.1"
```

```
## ggfun                "ggfun"                "0.0.9"
## ggnewscale           "ggnewscale"           "0.4.8"
## ggplot2              "ggplot2"              "3.4.0"
## ggplotify            "ggplotify"            "0.1.0"
## ggpubr               "ggpubr"               "0.5.0"
## ggraph               "ggraph"               "2.1.0"
## ggrepel              "ggrepel"              "0.9.2"
## ggsci                "ggsci"                "2.9"
## ggsignif             "ggsignif"             "0.6.4"
## ggstance             "ggstance"             "0.3.6"
## ggtree               "ggtree"               "3.6.2"
## gh                   "gh"                   "1.3.1"
## gitcreds             "gitcreds"             "0.1.2"
## glmnet               "glmnet"               "4.1-6"
## GlobalOptions        "GlobalOptions"        "0.1.2"
## globals              "globals"              "0.16.2"
## glue                 "glue"                 "1.6.2"
## GO.db                "GO.db"                "3.16.0"
## googledrive          "googledrive"          "2.0.0"
## googlesheets4        "googlesheets4"        "1.0.1"
## GOSemSim             "GOSemSim"             "2.24.0"
## GOstats              "GOstats"              "2.64.0"
## gower                "gower"                "1.0.1"
## GPfit                "GPfit"                "1.0-8"
## gplots               "gplots"               "3.1.3"
## gprofiler2           "gprofiler2"           "0.2.1"
## graph                "graph"                "1.76.0"
## graphlayouts         "graphlayouts"         "0.8.4"
## gridBase             "gridBase"             "0.4-7"
## gridExtra            "gridExtra"            "2.3"
## gridGraphics         "gridGraphics"         "0.5-1"
## GSEABase             "GSEABase"             "1.60.0"
## gson                 "gson"                 "0.0.9"
## gsubfn               "gsubfn"               "0.7"
## gtable               "gtable"               "0.3.1"
## gtools               "gtools"               "3.9.4"
## hardhat              "hardhat"              "1.2.0"
## hash                 "hash"                 "2.2.6.2"
## haven                "haven"                "2.5.1"
## HDF5Array            "HDF5Array"            "1.26.0"
## HDO.db               "HDO.db"               "0.99.1"
## here                 "here"                 "1.0.1"
## hexbin               "hexbin"               "1.28.2"
## highr                "highr"                "0.10"
## hms                  "hms"                  "1.1.2"
## htmltools            "htmltools"            "0.5.4"
## htmlwidgets          "htmlwidgets"          "1.6.0"
## httpuv               "httpuv"               "1.6.7"
## httr                 "httr"                 "1.4.4"
## hwriter              "hwriter"              "1.3.2.1"
## ids                  "ids"                  "1.0.1"
## igraph               "igraph"               "1.3.5"
## illuminaio           "illuminaio"           "0.40.0"
## infer                "infer"                "1.0.4"
## ini                  "ini"                  "0.3.1"
```

```
## insight               "insight"               "0.18.8"
## interactiveDisplayBase "interactiveDisplayBase" "1.36.0"
## inum                  "inum"                  "1.0-4"
## invgamma              "invgamma"              "1.1"
## ipred                 "ipred"                 "0.9-13"
## IRanges               "IRanges"               "2.32.0"
## IRdisplay             "IRdisplay"             "1.1"
## IRkernel              "IRkernel"              "1.3.1"
## irlba                 "irlba"                 "2.3.5.1"
## isoband               "isoband"               "0.2.7"
## iterators             "iterators"             "1.0.14"
## jquerylib             "jquerylib"             "0.1.4"
## jsonlite              "jsonlite"              "1.8.4"
## KEGGREST              "KEGGREST"              "1.38.0"
## kernlab               "kernlab"               "0.9-31"
## klaR                  "klaR"                  "1.7-1"
## knitr                 "knitr"                 "1.41"
## kohonen               "kohonen"               "3.0.11"
## labeling              "labeling"              "0.4.2"
## labelled              "labelled"              "2.10.0"
## lambda.r              "lambda.r"              "1.2.4"
## later                 "later"                 "1.3.0"
## lava                  "lava"                  "1.7.0"
## lazyeval              "lazyeval"              "0.2.2"
## leaps                 "leaps"                 "3.1"
## lhs                   "lhs"                   "1.1.6"
## libcoin               "libcoin"               "1.0-9"
## LiblineaR             "LiblineaR"             "2.10-22"
## lifecycle             "lifecycle"             "1.0.3"
## limma                 "limma"                 "3.54.0"
## listenv               "listenv"               "0.9.0"
## littler               "littler"               "0.3.17"
## lme4                  "lme4"                  "1.1-31"
## locfit                "locfit"                "1.5-9.7"
## lsa                   "lsa"                   "0.73.3"
## lubridate             "lubridate"             "1.9.0"
## magicaxis             "magicaxis"             "2.2.14"
## magrittr              "magrittr"              "2.0.3"
## mapproj               "mapproj"               "1.2.9"
## maps                  "maps"                  "3.4.1"
## margins               "margins"               "0.3.26"
## markdown              "markdown"              "1.4"
## marray                "marray"                "1.76.0"
## mashr                 "mashr"                 "0.2.69"
## MASS                  "MASS"                  "7.3-58.1"
## matrixcalc            "matrixcalc"            "1.0-6"
## MatrixGenerics        "MatrixGenerics"        "1.10.0"
## MatrixModels          "MatrixModels"          "0.5-1"
## matrixStats           "matrixStats"           "0.63.0"
## mclust                "mclust"                "6.0.0"
## memoise               "memoise"               "2.0.1"
## mime                  "mime"                  "0.12"
## minfi                 "minfi"                 "1.44.0"
## miniUI                "miniUI"                "0.1.1.1"
## minqa                 "minqa"                 "1.2.5"
```

```
## mixsqp                 "mixsqp"                 "0.3-48"
## modeldata              "modeldata"              "1.0.1"
## modelenv               "modelenv"               "0.1.0"
## ModelMetrics           "ModelMetrics"           "1.2.2.2"
## modelr                 "modelr"                 "0.1.10"
## multcompView           "multcompView"           "0.1-8"
## MultiAssayExperiment   "MultiAssayExperiment"   "1.24.0"
## multtest               "multtest"               "2.54.0"
## munsell                "munsell"                "0.5.0"
## mvtnorm                "mvtnorm"                "1.1-3"
## naivebayes             "naivebayes"             "0.9.7"
## netZooR                "netZooR"                "1.2.1"
## NISTunits              "NISTunits"              "1.0.1"
## nloptr                 "nloptr"                 "2.0.3"
## NMF                    "NMF"                    "0.25"
## nnet                   "nnet"                   "7.3-18"
## nor1mix                "nor1mix"                "1.3-0"
## numDeriv               "numDeriv"               "2016.8-1.1"
## openssl                "openssl"                "2.0.5"
## org.Hs.eg.db           "org.Hs.eg.db"           "3.16.0"
## pamr                   "pamr"                   "1.56.1"
## pandaR                 "pandaR"                 "1.30.0"
## parallelly             "parallelly"             "1.33.0"
## parameters             "parameters"             "0.20.0"
## parsnip                "parsnip"                "1.0.3"
## partykit               "partykit"               "1.2-16"
## pasilla                "pasilla"                "1.26.0"
## patchwork              "patchwork"              "1.1.2"
## pbdZMQ                 "pbdZMQ"                 "0.3-8"
## pbkrtest               "pbkrtest"               "0.5.1"
## penalized              "penalized"              "0.9-52"
## permute                "permute"                "0.9-7"
## PharmacoGx             "PharmacoGx"             "3.2.0"
## pheatmap               "pheatmap"               "1.0.12"
## piano                  "piano"                  "2.14.0"
## pillar                 "pillar"                 "1.8.1"
## pkgbuild               "pkgbuild"               "1.3.1"
## pkgconfig              "pkgconfig"              "2.0.3"
## pkgdown                "pkgdown"                "2.0.6"
## pkgload                "pkgload"                "1.3.1"
## PLIER                  "PLIER"                  "0.99.0"
## plogr                  "plogr"                  "0.2.0"
## plotly                 "plotly"                 "4.10.1"
## plotmo                 "plotmo"                 "3.6.2"
## plotrix                "plotrix"                "3.8-2"
## plyr                   "plyr"                   "1.8.8"
## png                    "png"                    "0.1-8"
## polyclip               "polyclip"               "1.10-4"
## polynom                "polynom"                "1.4-1"
## pracma                 "pracma"                 "2.4.2"
## praise                 "praise"                 "1.0.0"
## prediction             "prediction"             "0.3.14"
## preprocessCore         "preprocessCore"         "1.60.1"
## prettyunits            "prettyunits"            "1.1.1"
## pROC                   "pROC"                   "1.18.0"
```

```
## processx              "processx"              "3.8.0"
## prodlim               "prodlim"               "2019.11.13"
## profmem               "profmem"               "0.6.0"
## profvis               "profvis"               "0.3.7"
## progress              "progress"              "1.2.2"
## progressr             "progressr"             "0.12.0"
## projectR              "projectR"              "1.14.0"
## ProliferativeIndex    "ProliferativeIndex"    "1.0.1"
## promises              "promises"              "1.2.0.1"
## ProtGenerics          "ProtGenerics"          "1.30.0"
## proto                 "proto"                 "1.0.0"
## proxy                 "proxy"                 "0.4-27"
## ps                    "ps"                    "1.7.2"
## purrr                 "purrr"                 "1.0.0"
## quadprog              "quadprog"              "1.5-8"
## quantreg              "quantreg"              "5.94"
## quantro               "quantro"               "1.32.0"
## questionr             "questionr"             "0.7.7"
## qvalue                "qvalue"                "2.30.0"
## R.cache               "R.cache"               "0.16.0"
## R.methodsS3           "R.methodsS3"           "1.8.2"
## R.oo                  "R.oo"                  "1.25.0"
## R.utils               "R.utils"               "2.12.2"
## R6                    "R6"                    "2.5.1"
## ragg                  "ragg"                  "1.2.4"
## randomForest          "randomForest"          "4.7-1.1"
## ranger                "ranger"                "0.14.1"
## RANN                  "RANN"                  "2.6.1"
## rappdirs              "rappdirs"              "0.3.3"
## RBGL                  "RBGL"                  "1.74.0"
## rcmdcheck             "rcmdcheck"             "1.4.0"
## RColorBrewer          "RColorBrewer"          "1.1-3"
## Rcpp                  "Rcpp"                  "1.0.9"
## RcppArmadillo         "RcppArmadillo"         "0.11.4.2.1"
## RcppEigen             "RcppEigen"             "0.3.3.9.3"
## RcppGSL               "RcppGSL"               "0.3.12"
## RcppTOML              "RcppTOML"              "0.1.7"
## RCurl                 "RCurl"                 "1.98-1.9"
## RCy3                  "RCy3"                  "2.18.0"
## reactome.db           "reactome.db"           "1.82.0"
## readr                 "readr"                 "2.1.3"
## readxl                "readxl"                "1.4.1"
## recipes               "recipes"               "1.0.3"
## recount3              "recount3"              "1.8.0"
## registry              "registry"              "0.5-1"
## relations             "relations"             "0.6-12"
## rematch               "rematch"               "1.0.1"
## rematch2              "rematch2"              "2.1.2"
## remotes               "remotes"               "2.4.2"
## repr                  "repr"                  "1.1.4"
## reprex                "reprex"                "2.0.2"
## reshape               "reshape"               "0.8.9"
## reshape2              "reshape2"              "1.4.4"
## restfulr              "restfulr"              "0.0.15"
## reticulate            "reticulate"            "1.26"
```

```
## Rgraphviz              "Rgraphviz"              "2.42.0"
## rhdf5                  "rhdf5"                  "2.42.0"
## rhdf5filters           "rhdf5filters"           "1.10.0"
## Rhdf5lib               "Rhdf5lib"               "1.20.0"
## Rhtslib                "Rhtslib"                "2.0.0"
## rJava                  "rJava"                  "1.0-6"
## rjson                  "rjson"                  "0.2.21"
## RJSONIO                "RJSONIO"                "1.3-1.6"
## rlang                  "rlang"                  "1.0.6"
## rmarkdown              "rmarkdown"              "2.19"
## rmeta                  "rmeta"                  "3.0"
## rngtools               "rngtools"               "1.5.2"
## robustbase             "robustbase"             "0.95-0"
## ROCR                   "ROCR"                   "1.0-11"
## roxygen2               "roxygen2"               "7.2.1"
## rpart                  "rpart"                  "4.1.19"
## rprojroot              "rprojroot"              "2.0.3"
## rsample                "rsample"                "1.1.1"
## Rsamtools              "Rsamtools"              "2.14.0"
## RSQLite                "RSQLite"                "2.2.20"
## rstatix                "rstatix"                "0.7.1"
## rstudioapi             "rstudioapi"             "0.14"
## rsvd                   "rsvd"                   "1.0.5"
## rsvg                   "rsvg"                   "2.4.0"
## rtracklayer            "rtracklayer"            "1.58.0"
## RUnit                  "RUnit"                  "0.4.32"
## rversions              "rversions"              "2.1.2"
## rvest                  "rvest"                  "1.0.3"
## RWeka                  "RWeka"                  "0.4-44"
## RWekajars              "RWekajars"              "3.9.3-2"
## S4Vectors              "S4Vectors"              "0.36.1"
## sass                   "sass"                   "0.4.4"
## scales                 "scales"                 "1.2.1"
## scatterpie             "scatterpie"             "0.1.8"
## scatterplot3d          "scatterplot3d"          "0.3-42"
## scrime                 "scrime"                 "1.3.5"
## selectr                "selectr"                "0.4-2"
## sessioninfo            "sessioninfo"            "1.2.2"
## sets                   "sets"                   "1.0-21"
## shadowtext             "shadowtext"             "0.1.2"
## shape                  "shape"                  "1.4.6"
## shiny                  "shiny"                  "1.7.4"
## shinydashboard         "shinydashboard"         "0.7.2"
## shinyjs                "shinyjs"                "2.1.0"
## siggenes               "siggenes"               "1.72.0"
## signatureSearch        "signatureSearch"        "1.11.1"
## signatureSearchData    "signatureSearchData"    "1.12.0"
## SingleCellExperiment   "SingleCellExperiment"   "1.20.0"
## skimr                  "skimr"                  "2.1.5"
## slam                   "slam"                   "0.1-50"
## slider                 "slider"                 "0.3.0"
## sm                     "sm"                     "2.2-5.7.1"
## snow                   "snow"                   "0.4-4"
## SnowballC              "SnowballC"              "0.7.0"
## softImpute             "softImpute"             "1.4-1"
```

```
## sourcetools           "sourcetools"           "0.1.7"
## SparseM               "SparseM"               "1.81"
## sparseMatrixStats     "sparseMatrixStats"     "1.10.0"
## sqldf                 "sqldf"                 "0.4-11"
## SQUAREM               "SQUAREM"               "2021.1"
## statmod               "statmod"               "1.4.37"
## STRINGdb              "STRINGdb"              "2.10.0"
## stringi               "stringi"               "1.7.8"
## stringr               "stringr"               "1.5.0"
## styler                "styler"                "1.8.1"
## SummarizedExperiment  "SummarizedExperiment"  "1.28.0"
## sys                   "sys"                   "3.4.1"
## systemfonts           "systemfonts"           "1.0.4"
## TeachingDemos         "TeachingDemos"         "2.12"
## testthat              "testthat"              "3.1.5"
## textshaping           "textshaping"           "0.3.6"
## TFEA.ChIP             "TFEA.ChIP"             "1.18.0"
## tibble                "tibble"                "3.1.8"
## tidygraph             "tidygraph"             "1.2.2"
## tidymodels            "tidymodels"            "1.0.0"
## tidyr                 "tidyr"                 "1.2.1"
## tidyselect            "tidyselect"            "1.2.0"
## tidytree              "tidytree"              "0.4.2"
## tidyverse             "tidyverse"             "1.3.2"
## timechange            "timechange"            "0.1.1"
## timeDate              "timeDate"              "4021.107"
## tinytex               "tinytex"               "0.43"
## topGO                 "topGO"                 "2.50.0"
## treeio                "treeio"                "1.22.0"
## truncnorm             "truncnorm"             "1.0-8"
## tune                  "tune"                  "1.0.1"
## tweenr                "tweenr"                "2.0.2"
## tzdb                  "tzdb"                  "0.3.0"
## uchardet              "uchardet"              "1.1.1"
## urlchecker            "urlchecker"            "1.0.1"
## usethis               "usethis"               "2.1.6"
## utf8                  "utf8"                  "1.2.2"
## uuid                  "uuid"                  "1.1-0"
## vctrs                 "vctrs"                 "0.5.1"
## vegan                 "vegan"                 "2.6-4"
## VennDiagram           "VennDiagram"           "1.7.3"
## viridis               "viridis"               "0.6.2"
## viridisLite           "viridisLite"           "0.4.1"
## visNetwork            "visNetwork"            "2.1.2"
## vroom                 "vroom"                 "1.6.0"
## waldo                 "waldo"                 "0.4.0"
## warp                  "warp"                  "0.2.0"
## whisker               "whisker"               "0.4"
## withr                 "withr"                 "2.5.0"
## workflows             "workflows"             "1.1.2"
## workflowsets          "workflowsets"          "1.0.0"
## xfun                  "xfun"                  "0.36"
## xgboost               "xgboost"               "1.6.0.1"
## XML                   "XML"                   "3.99-0.13"
## xml2                  "xml2"                  "1.3.3"
```

```
## xopen                "xopen"               "1.0.0"
## xtable               "xtable"              "1.8-4"
## XVector              "XVector"             "0.38.0"
## yaml                 "yaml"                "2.3.6"
## yardstick            "yardstick"           "1.1.0"
## yarn                 "yarn"                "1.24.0"
## yulab.utils          "yulab.utils"         "0.0.6"
## zip                  "zip"                 "2.2.2"
## zlibbioc             "zlibbioc"            "1.44.0"
## base                 "base"                "4.2.2"
## boot                 "boot"                "1.3-28"
## class                "class"               "7.3-20"
## cluster              "cluster"             "2.1.4"
## codetools            "codetools"           "0.2-18"
## compiler             "compiler"            "4.2.2"
## datasets             "datasets"            "4.2.2"
## foreign              "foreign"             "0.8-83"
## graphics             "graphics"            "4.2.2"
## grDevices            "grDevices"           "4.2.2"
## grid                 "grid"                "4.2.2"
## KernSmooth           "KernSmooth"          "2.23-20"
## lattice              "lattice"             "0.20-45"
## MASS                 "MASS"                "7.3-58.1"
## Matrix               "Matrix"              "1.5-1"
## methods              "methods"             "4.2.2"
## mgcv                 "mgcv"                "1.8-41"
## nlme                 "nlme"                "3.1-160"
## nnet                 "nnet"                "7.3-18"
## parallel             "parallel"            "4.2.2"
## rpart                "rpart"               "4.1.19"
## spatial              "spatial"             "7.3-15"
## splines              "splines"             "4.2.2"
## stats                "stats"               "4.2.2"
## stats4               "stats4"              "4.2.2"
## survival             "survival"            "3.4-0"
## tcltk                "tcltk"               "4.2.2"
## tools                "tools"               "4.2.2"
## utils                "utils"               "4.2.2"
```

```
R version 4.0.5 (2021-03-31)
Platform: x86_64-conda-linux-gnu (64-bit)
Running under: Red Hat Enterprise Linux

Matrix products: default
BLAS/LAPACK: /data/user/jfisher7/.conda/envs/SR_TAU_CELL/lib/
libopenblasp-r0.3.18.so

locale:
[1] C

attached base packages:
[1] stats     graphics  grDevices utils     datasets  methods   base

loaded via a namespace (and not attached):
[1] compiler_4.0.5
[1] "Listing packages"
                       Package                  Version
AnnotationDbi          "AnnotationDbi"          "1.52.0"
AnnotationHub          "AnnotationHub"          "2.22.1"
BH                     "BH"                     "1.78.0-0"
Biobase                "Biobase"                "2.50.0"
BiocFileCache          "BiocFileCache"          "1.14.0"
BiocGenerics           "BiocGenerics"           "0.36.1"
BiocManager            "BiocManager"            "1.30.16"
BiocParallel           "BiocParallel"           "1.24.1"
BiocVersion            "BiocVersion"            "3.12.0"
DBI                    "DBI"                    "1.1.2"
DO.db                  "DO.db"                  "2.9"
DOSE                   "DOSE"                   "3.16.0"
DT                     "DT"                     "0.20"
DelayedArray           "DelayedArray"           "0.16.3"
ExperimentHub          "ExperimentHub"          "1.16.1"
GO.db                  "GO.db"                  "3.12.1"
GOSemSim               "GOSemSim"               "2.16.1"
GSEABase               "GSEABase"               "1.52.1"
GenomeInfoDb           "GenomeInfoDb"           "1.26.7"
GenomeInfoDbData       "GenomeInfoDbData"       "1.2.4"
GenomicRanges          "GenomicRanges"          "1.42.0"
HDF5Array              "HDF5Array"              "1.18.1"
IRanges                "IRanges"                "2.24.1"
MASS                   "MASS"                   "7.3-55"
Matrix                 "Matrix"                 "1.4-0"
MatrixGenerics         "MatrixGenerics"         "1.2.1"
R.methodsS3            "R.methodsS3"            "1.8.1"
R.oo                   "R.oo"                   "1.24.0"
R.utils                "R.utils"                "2.11.0"
R6                     "R6"                     "2.5.1"
RColorBrewer           "RColorBrewer"           "1.1-2"
RCurl                  "RCurl"                  "1.98-1.5"
```

```
RSQLite                 "RSQLite"                "2.2.9"
Rcpp                    "Rcpp"                   "1.0.8"
RcppArmadillo           "RcppArmadillo"          "0.10.8.1.0"
RcppEigen               "RcppEigen"              "0.3.3.9.1"
Rhdf5lib                "Rhdf5lib"               "1.12.1"
S4Vectors               "S4Vectors"              "0.28.1"
SummarizedExperiment    "SummarizedExperiment"   "1.20.0"
XML                     "XML"                    "3.99-0.8"
XVector                 "XVector"                "0.30.0"
affy                    "affy"                   "1.68.0"
affyio                  "affyio"                 "1.60.0"
annotate                "annotate"               "1.68.0"
askpass                 "askpass"                "1.1"
assertthat              "assertthat"             "0.2.1"
backports               "backports"              "1.4.1"
base                    "base"                   "4.0.5"
base64enc               "base64enc"              "0.1-3"
bit                     "bit"                    "4.0.4"
bit64                   "bit64"                  "4.0.5"
bitops                  "bitops"                 "1.0-7"
blob                    "blob"                   "1.2.2"
brio                    "brio"                   "1.1.3"
broom                   "broom"                  "0.7.12"
bslib                   "bslib"                  "0.3.1"
cachem                  "cachem"                 "1.0.6"
callr                   "callr"                  "3.7.0"
cellranger              "cellranger"             "1.1.0"
cli                     "cli"                    "3.1.1"
clipr                   "clipr"                  "0.7.1"
clusterProfiler         "clusterProfiler"        "3.18.1"
colorspace              "colorspace"             "2.0-2"
commonmark              "commonmark"             "1.7"
compiler                "compiler"               "4.0.5"
cowplot                 "cowplot"                "1.1.1"
cpp11                   "cpp11"                  "0.4.2"
crayon                  "crayon"                 "1.4.2"
crosstalk               "crosstalk"              "1.2.0"
curl                    "curl"                   "4.3.2"
data.table              "data.table"             "1.14.2"
datasets                "datasets"               "4.0.5"
dbplyr                  "dbplyr"                 "2.1.1"
desc                    "desc"                   "1.4.0"
diffobj                 "diffobj"                "0.3.5"
digest                  "digest"                 "0.6.29"
downloader              "downloader"             "0.4"
dplyr                   "dplyr"                  "1.0.7"
dtplyr                  "dtplyr"                 "1.2.1"
ellipsis                "ellipsis"               "0.3.2"
enrichplot              "enrichplot"             "1.10.2"
evaluate                "evaluate"               "0.14"
```

| | | |
|---|---|---|
| fansi | "fansi" | "1.0.2" |
| farver | "farver" | "2.1.0" |
| fastmap | "fastmap" | "1.1.0" |
| fastmatch | "fastmatch" | "1.1-3" |
| fgsea | "fgsea" | "1.16.0" |
| fontawesome | "fontawesome" | "0.2.2" |
| forcats | "forcats" | "0.5.1" |
| formatR | "formatR" | "1.11" |
| fs | "fs" | "1.5.2" |
| futile.logger | "futile.logger" | "1.4.3" |
| futile.options | "futile.options" | "1.0.1" |
| gargle | "gargle" | "1.2.0" |
| generics | "generics" | "0.1.2" |
| ggforce | "ggforce" | "0.3.3" |
| ggfun | "ggfun" | "0.0.5" |
| ggplot2 | "ggplot2" | "3.3.5" |
| ggraph | "ggraph" | "2.0.5" |
| ggrepel | "ggrepel" | "0.9.1" |
| glue | "glue" | "1.6.1" |
| googledrive | "googledrive" | "2.0.0" |
| googlesheets4 | "googlesheets4" | "1.0.0" |
| grDevices | "grDevices" | "4.0.5" |
| graph | "graph" | "1.68.0" |
| graphics | "graphics" | "4.0.5" |
| graphlayouts | "graphlayouts" | "0.8.0" |
| grid | "grid" | "4.0.5" |
| gridExtra | "gridExtra" | "2.3" |
| gtable | "gtable" | "0.3.0" |
| haven | "haven" | "2.4.3" |
| highr | "highr" | "0.9" |
| hms | "hms" | "1.1.1" |
| htmltools | "htmltools" | "0.5.2" |
| htmlwidgets | "htmlwidgets" | "1.5.4" |
| httpuv | "httpuv" | "1.6.5" |
| httr | "httr" | "1.4.2" |
| ids | "ids" | "1.0.1" |
| igraph | "igraph" | "1.2.11" |
| interactiveDisplayBase | "interactiveDisplayBase" | "1.28.0" |
| isoband | "isoband" | "0.2.5" |
| jquerylib | "jquerylib" | "0.1.4" |
| jsonlite | "jsonlite" | "1.7.3" |
| knitr | "knitr" | "1.37" |
| labeling | "labeling" | "0.4.2" |
| lambda.r | "lambda.r" | "1.2.4" |
| later | "later" | "1.3.0" |
| lattice | "lattice" | "0.20-45" |
| lazyeval | "lazyeval" | "0.2.2" |
| lifecycle | "lifecycle" | "1.0.1" |
| limma | "limma" | "3.46.0" |
| lubridate | "lubridate" | "1.8.0" |

```
magrittr              "magrittr"              "2.0.2"
matrixStats           "matrixStats"           "0.61.0"
memoise               "memoise"               "2.0.1"
methods               "methods"               "4.0.5"
mgcv                  "mgcv"                  "1.8-38"
mime                  "mime"                  "0.12"
modelr                "modelr"                "0.1.8"
munsell               "munsell"               "0.5.0"
nlme                  "nlme"                  "3.1-155"
openssl               "openssl"               "1.4.6"
parallel              "parallel"              "4.0.5"
pillar                "pillar"                "1.7.0"
pkgconfig             "pkgconfig"             "2.0.3"
pkgload               "pkgload"               "1.2.4"
plogr                 "plogr"                 "0.2.0"
plyr                  "plyr"                  "1.8.6"
polyclip              "polyclip"              "1.10-0"
praise                "praise"                "1.0.0"
preprocessCore        "preprocessCore"        "1.52.1"
prettyunits           "prettyunits"           "1.1.1"
processx              "processx"              "3.5.2"
progress              "progress"              "1.2.2"
promises              "promises"              "1.2.0.1"
ps                    "ps"                    "1.6.0"
purrr                 "purrr"                 "0.3.4"
qvalue                "qvalue"                "2.22.0"
rappdirs              "rappdirs"              "0.3.3"
reactome.db           "reactome.db"           "1.74.0"
readr                 "readr"                 "2.1.2"
readxl                "readxl"                "1.3.1"
rematch               "rematch"               "1.0.1"
rematch2              "rematch2"              "2.1.2"
reprex                "reprex"                "2.0.1"
reshape2              "reshape2"              "1.4.4"
rhdf5                 "rhdf5"                 "2.34.0"
rhdf5filters          "rhdf5filters"          "1.2.1"
rlang                 "rlang"                 "1.0.0"
rmarkdown             "rmarkdown"             "2.11"
rprojroot             "rprojroot"             "2.0.2"
rstudioapi            "rstudioapi"            "0.13"
rvcheck               "rvcheck"               "0.2.1"
rvest                 "rvest"                 "1.0.2"
sass                  "sass"                  "0.4.0"
scales                "scales"                "1.1.1"
scatterpie            "scatterpie"            "0.1.7"
selectr               "selectr"               "0.4-2"
shadowtext            "shadowtext"            "0.1.1"
shiny                 "shiny"                 "1.7.1"
signatureSearch       "signatureSearch"       "1.4.6"
signatureSearchData   "signatureSearchData"   "1.4.0"
```

| | | |
|---|---|---|
| snow | "snow" | "0.4-4" |
| sourcetools | "sourcetools" | "0.1.7" |
| splines | "splines" | "4.0.5" |
| stats | "stats" | "4.0.5" |
| stats4 | "stats4" | "4.0.5" |
| stringi | "stringi" | "1.7.6" |
| stringr | "stringr" | "1.4.0" |
| sys | "sys" | "3.4" |
| tcltk | "tcltk" | "4.0.5" |
| testthat | "testthat" | "3.1.2" |
| tibble | "tibble" | "3.1.6" |
| tidygraph | "tidygraph" | "1.2.0" |
| tidyr | "tidyr" | "1.2.0" |
| tidyselect | "tidyselect" | "1.1.1" |
| tidyverse | "tidyverse" | "1.3.1" |
| tinytex | "tinytex" | "0.36" |
| tools | "tools" | "4.0.5" |
| tweenr | "tweenr" | "1.0.2" |
| tzdb | "tzdb" | "0.2.0" |
| utf8 | "utf8" | "1.2.2" |
| utils | "utils" | "4.0.5" |
| uuid | "uuid" | "1.0-3" |
| vctrs | "vctrs" | "0.3.8" |
| viridis | "viridis" | "0.6.2" |
| viridisLite | "viridisLite" | "0.4.0" |
| visNetwork | "visNetwork" | "2.1.0" |
| vroom | "vroom" | "1.5.7" |
| waldo | "waldo" | "0.3.1" |
| withr | "withr" | "2.4.3" |
| xfun | "xfun" | "0.29" |
| xml2 | "xml2" | "1.3.3" |
| xtable | "xtable" | "1.8-4" |
| yaml | "yaml" | "2.2.2" |
| yulab.utils | "yulab.utils" | "0.0.4" |
| zlibbioc | "zlibbioc" | "1.36.0" |
